# SNARE assembly enlightened by cryo-EM structures of a synaptobrevin-Munc18-1-syntaxin-1 complex

**DOI:** 10.1101/2022.03.05.483126

**Authors:** Karolina P. Stepien, Junjie Xu, Xuewu Zhang, Xiaochen Bai, Josep Rizo

## Abstract

Munc18-1 forms a template to recruit and organize assembly of the SNARE complex that triggers neurotransmitter release, binding first to a closed conformation of syntaxin-1 where its N-terminal region interacts with the SNARE motif, and later binding to synaptobrevin. However, the mechanism of SNARE complex assembly remains unclear. Here we report two cryo-EM structures of Munc18-1 bound to cross-linked syntaxin-1 and synaptobrevin. The structures allow visualization of how syntaxin-1 opens and reveal how part of the syntaxin-1 N-terminal region can help nucleate interactions between the N-termini of the syntaxin-1 and synaptobrevin SNARE motifs while their C-termini bind to distal sites of Munc18-1. Mutagenesis, SNARE complex assembly assays and reconstitution experiments support a model whereby these interactions are critical to initiate SNARE complex assembly.

**One Sentence Summary:** Cryo-EM structures reveal key insights into the molecular mechanism of neuronal SNARE complex assembly templated by Munc18-1

## Main Text

Release of neurotransmitters by Ca^2+^-evoked synaptic vesicle exocytosis is mediated by the vesicle soluble N-ethylmaleimide-sensitive factor attachment protein receptor (SNARE) synaptobrevin and the plasma membrane SNAREs syntaxin-1 and SNAP-25, which form a tight complex (*1*) through their ∼65-residue SNARE motifs. The SNARE complex consists of a parallel four-helix bundle (*2, 3*) and brings the vesicle and plasma membranes together (*4*), which is crucial for membrane fusion [reviewed in (*5*)]. In syntaxin-1, the SNARE motif is preceded by a 190-residue N-terminal region (Fig. 1A) containing a three-helix bundle domain (the H_abc_ domain) that binds intramolecularly to the SNARE motif, forming a ‘closed’ conformation that precludes SNARE complex assembly (*6-8*). The SNARE complex is disassembled by N-ethylmaleimide-sensitive factor (NSF) and soluble NSF attachment proteins (SNAPs; no relation to SNAP-25) (*1*), whereas SNARE complex assembly is orchestrated through an NSF-SNAP-resistant pathway by Munc18-1 and Munc13 (*9, 10*). This pathway starts with Munc18-1 bound tightly to closed syntaxin-1 (*7, 8*) and is activated by Munc13 when it helps to open syntaxin-1 (*11, 12*) while it bridges the vesicle and plasma membranes (*13, 14*), enabling a wide variety of Munc13-dependent presynaptic plasticity processes that underlie multiple forms of information processing in the brain (*15-17*). The importance of this pathway was highlighted by the total abrogation of neurotransmitter release observed in the absence of Munc18-1 or Munc13s (*18-21*). Moreover, an L165A,E166A mutation that opens syntaxin-1 (LE mutation) (*7*) partially rescues the impairments in release caused by deletion of diverse proteins in *C. elegans*, including the Munc13 homologue Unc-13 (*15, 22, 23*), and a P335A mutation in Munc18-1 leads to a gain-of function (*24, 25*) that also rescues release partially in *unc-13* nulls (*16*). Thus, elucidating the mechanism of SNARE complex assembly is not only critical to understand brain function but can help develop novel therapies for many neurological disorders exhibiting altered synaptic transmission, and is also relevant to diseases arising from defects in regulated secretion, including hypertension, cancer and diabetes (*26, 27*).

**Fig. 1.**
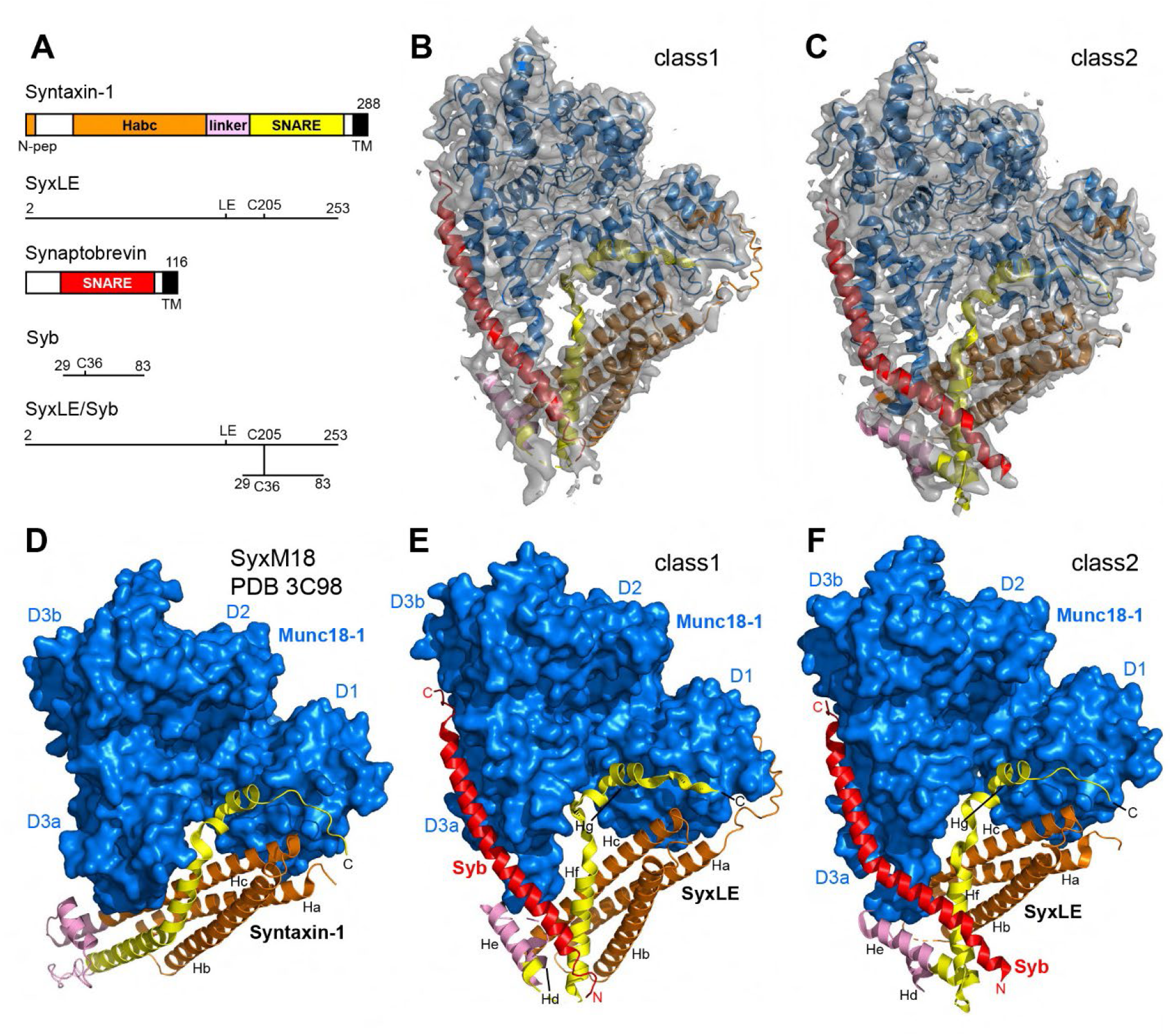
Two cryo-EM structures of the template complex. (**A**) Domain diagrams of syntaxin-1 and synaptobrevin, and summary of the fragments used to prepare SyxLE/Syb. N-pep = N-peptide. SNARE = SNARE motif. (**B**,**C**) 3D reconstructions of two structures of the template complex, class1 (**A**) and class2 (**B**), and corresponding ribbon diagrams fitted into the cryo-EM maps at 3.7 and 3.5 Å, respectively. (**D-F**) Comparison of the crystal structure of the syntaxin-1-Munc18-1 complex (SyxM18) (PDB code 3C98) (*8, 32*) (**D**) with the two cryo-EM structures of the template complex, class1 (**E**) and class2 (**F**). The surface of Munc18-1 is shown in blue and the SNAREs are represented by ribbon diagrams with synaptobrevin (Syb) in red and syntaxin-1 in orange (N-peptide and H_abc_ domain), pink (linker) and yellow (SNARE motif). The domains of Munc18-1 (D1, D2, D3a and D3b) are labeled. The helices formed by syntaxin-1 in class1 and class2 (named Ha-Hg) are indicated. Helices Ha-Hc are also present in the syntaxin-1-Munc18-1 complex, but the other helices are remodeled. The N- and C-termini of Syb, as well as the C-terminus of SyxLE are labeled.

In addition to binding tightly to closed syntaxin-1, Munc18-1 interacts weakly with synaptobrevin, which suggested that Munc18-1 forms a template for SNARE complex assembly (*24*). This notion was firmly established by seminal crystal structures of the yeast vacuolar Sec1-Munc18 (SM) homologue Vps33 bound to the SNARE motifs of the syntaxin-1 homologue Vam3 or the synaptobrevin homologue Nyv1 (*28*), which showed that simultaneous binding to Vps33 would place Vam3 and Nyv1 in the correct register for SNARE complex assembly. Subsequent biophysical studies of Munc18-1 further supported the idea that a template complex of Munc18-1 bound to syntaxin-1 and synaptobrevin is a central intermediate between the Munc18-1-closed syntaxin-1 complex and the SNARE complex, and showed that the syntaxin-1 N-terminal region plays a crucial role in template complex formation (*29, 30*). However, the basis for such crucial role is unknown, and the steps that lead from the Munc18-1-closed syntaxin-1 complex to the SNARE complex remain enigmatic, in part because no structures are available for Munc18-1 bound to synaptobrevin or for any SM protein bound simultaneously to its two cognate SNAREs.

Here we fill this fundamental gap, describing the elucidation of two structures of a synaptobrevin-Munc18-1-syntaxin-1 template complex by cryo-electron microscopy (cryo-EM). The structures reveal how Munc18-1 holds synaptobrevin and syntaxin-1 through the C-termini of their SNARE motifs while the syntaxin-1 conformation opens gradually and the syntaxin-1 N-terminal region helps to nucleate SNARE assembly, explaining the critical role of this region.

### Cryo-EM structures of a template complex

The weak affinity of synaptobrevin for Munc18-1 (*31*) hinders structural studies of complexes between them. To overcome this problem, we cross-linked the syntaxin-1 cytoplasmic region containing the LE mutation (SyxLE) to a synaptobrevin fragment spanning most of its SNARE motif (Syb) through a disulfilde bond between single cysteine residues placed at the N-termini of their SNARE motifs (SyxLE/Syb; Fig. 1A). This approach was shown to facilitate formation of a synaptobrevin-Munc18-1-syntaxin-1 template complex (*30*). To further favor template complex formation, we used Munc18-1 bearing a D326K gain-of-function mutation that increases the affinity of synaptobrevin for Munc18-1 because it unfurls a Munc18-1 loop that covers the synaptobrevin binding site (*29*). Indeed, SyxLE/Syb co-eluted with Munc18-1 D326K in gel filtration (Fig. S1A,B) and formed an SDS-resistant SNARE complex with SNAP-25 in the presence of Munc18-1 much more efficiently than the separate SyxLE and Syb fragments (Fig. S1C).

Imaging of the template complex by cryo-EM and initial 3D classification led to the identification of two major classes, which we refer to as class1 and class2 and were resolved at 3.7 and 3.5 Å, respectively (Figs. 1B,C, S2, S3, Table S1). The two structures reveal extensive interactions of Munc18-1 with syntaxin-1 and synaptobrevin (see buried surface areas in Table S2) and have similar architectures, with common features that were also observed in the crystal structure of the syntaxin-1-Munc18-1 complex (*8*) but with important differences (Figs. 1D-F, S4). Munc18-1 has an almost identical three-domain arch shape in the three structures. The most notable difference is in the loop joining helices H11 and H12 of domain 3a, which is furled in the Munc18-1-syntaxin-1 complex, covering the synaptobrevin binding site, but is unfurled in class1 and class2 to allow synaptobrevin binding (Fig. 2A-C). Almost the entire synaptobrevin SNARE motif forms a continuous helix in both class1 and class2, but in class2 the helix is bent at the middle (Fig. 1E,F). The C-terminal half of the synaptobrevin SNARE motif binds to a groove formed by helices H11 and H12 of Munc18-1 (Figs. 1E,F, 2B,C), in an analogous position to that observed for Nyv1 bound to Vps33 (*28*) (Fig. S5A-C). Binding to the groove is mediated by a combination of hydrophobic and polar interactions, and side chains in the interface include L307 and L348 of Munc18-1 (Fig. S6A-D), which had been implicated in synaptobrevin binding and/or template complex formation through mutagenesis (*24, 30*). The R56 side chain in the middle of the synaptobrevin SNARE motif binds to a pocket at the bottom of the H11-H12 groove in both class1 and class2, but there are almost no further contacts between Munc18-1 and residues N-terminal to R56 (Fig. S6E-H). This observation contrasts with the extensive contacts of the N-terminal half of Nyv1 with Vps33 in their complex (Fig. S5A). Instead, the

**Fig. 2.**
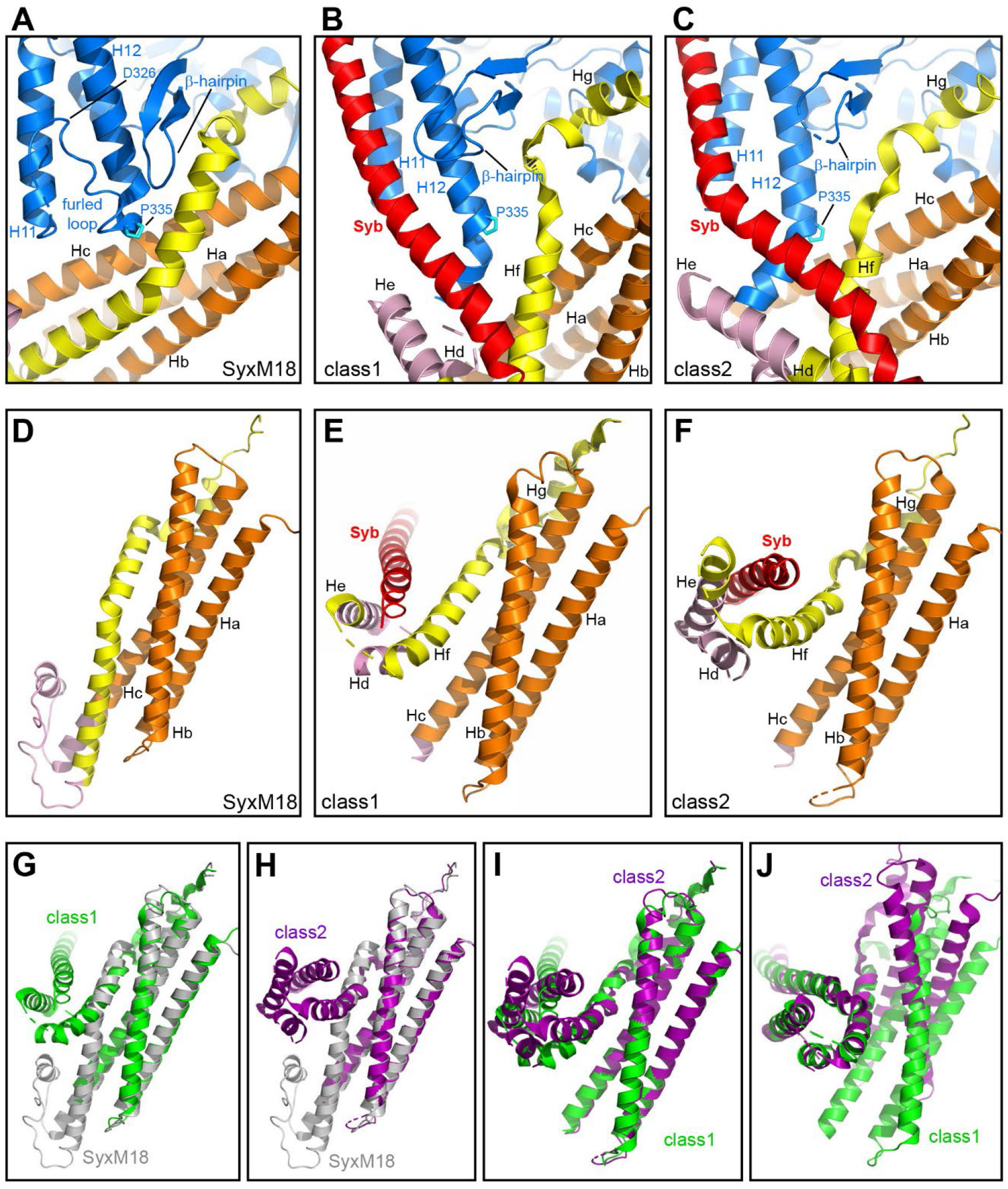
Structural changes leading to template complex formation. (**A**-**C**) Close-up views of the area where P335 binds to closed syntaxin-1 in the syntaxin-1-Munc18-1 complex (SyxM18) (**A**) and synaptobrevin binds to Munc18-1 and syntaxin-1 in class1 (**B**) and class2 (**C**). Munc18-1 is colored in blue, synaptobrevin (Syb) in red and syntaxin-1 in orange (N-peptide and H_abc_ domain), pink (linker) and yellow (SNARE motif). P335 is shown as a stick model with carbon atoms in cyan. The positions of the furled loop and D326 in the Munc18-1-syntaxin-1 complex, as well as of synaptobrevin (Syb), P335, a nearby β-hairpin and selected helices of Munc18-1 and syntaxin-1 are indicated. (**D**-**F**) Close-up views of the region where the syntaxin-1 SNARE motif interacts with the H_abc_ domain and the linker in SyxM18 (**D**) as well as with synaptobrevin in class1 (**E**) and class2 (**F**). Syb and helices Hd-Hf of syntaxin-1 form a small four-helix bundle in class1 and class2. (**G**-**I**) Superpositions among SyxM18 (grey), class1 (green) and class2 (purple) in the same close-up views shown in (**D**-**F**). The structures were superimposed using the atoms of the H_abc_ domain. (**J**) Superposition of class1 and class2 in a similar close-up view but superimposing the atoms of parts of the syntaxin-1 and synaptobrevin SNARE motifs (residues 206-219 and 37-50, respectively) to show that the small four-helix bundles that they form with helices Hd and He of the syntaxin-1 linker are similar in class1 and class2. Munc18-1 is not shown for simplicity in (**D-J**).

N-terminal half of the synaptobrevin SNARE motif interacts with syntaxin-1 (Figs. 1E,F, S6E-H), which is expected to be favored by the disulfide bond. It is noteworthy that R56 of synaptobrevin forms a polar layer in the middle of the SNARE complex four-helix bundle, which otherwise is formed by hydrophobic layers (*3*), and is at the corner where the synaptobrevin helix bends in class2 (Fig. S6H). Thus, R56 may constitute a pivot point where helices formed by the N-and C-terminal halves of the synaptobrevin SNARE motif may change direction during SNARE assembly.

In both class1 and class2, syntaxin-1 forms a short α-helix at the very N-terminus (the N-peptide motif) that binds to the N-terminal domain of Munc18-1, as in the syntaxin-1-Munc18-1 complex (*32*), and seven additional helices (called Ha-Hg) (Figs. 1E,F, S4B,C). The C-terminus of the syntaxin-1 SNARE motif, including the Hg helix, binds to a groove of domain 1 of Munc18-1 and a pocket that defines its arch shape, also as observed in the Munc18-1-syntaxin-1 complex (Figs. 1D-F, S7A-C). This interaction and that of the N-peptide are the most conserved among the multiple interactions between syntaxin-1 and Munc18-1 in their binary complex, class1 and class2, suggesting that they serve as anchor points that retain syntaxin-1 bound to Munc18-1 while other regions undergo substantial rearrangements in the pathway toward SNARE complex assembly. The structure of the H_abc_ domain remains largely unaltered in the three complexes and the H_abc_ domain-Munc18-1 interface is similar in the three structures (Figs. S7D-F), but the orientation of H_abc_ with respect to Munc18-1 is somewhat different in class1 (Fig. S4D-F), suggesting that this interface can adapt to structural rearrangements.

The N-terminal half of the syntaxin-1 SNARE motif forms a long α-helix (Hf) that contacts domain 3a of Munc18-1 in both class1 and class2 (Fig. 1E,F), and is in an analogous position to that of the homologous region of Vam3 in its complex with Vps33 (Fig. S5D-F). This region of syntaxin-1 also binds to the same site of domain 3a of Munc18-1 in the binary complex, but establishing more extensive contacts than in class1 and class2 (Figs. 2A-C, S6I-K). Moreover, the helix formed by this region is shifted and rotated in class1 and class2 with respect to its position in the Munc18-1-syntaxin-1 complex (Fig. S6I-K), where the helix is extended to the very N-terminus of the SNARE motif, exhibiting a bend in the middle (Fig. 1D) that is not observed in helix Hf of class1 and class2 (Fig. 1E,F). This feature of the Munc18-1-syntaxin-1 complex allows extensive interactions of the SNARE motif with the H_abc_ domain that define the closed conformation of syntaxin-1, but these contacts are less extensive in class1 and even less extensive in class2 (Table S2), reflecting a gradual rotation of the SNARE motif with respect to the H_abc_ domain that may reflect how the syntaxin-1 conformation opens (Figs. 2D-I, S7H-J). These observations suggest that class1 occurs first and class2 later.

In class1 and class2, the very N-terminus of the syntaxin-1 SNARE motif forms helix He together with part of the linker sequence connecting the H_abc_ domain with the SNARE motif (Figs. 1E,F, 2E,F). The linker region forms an additional helix (Hd), and adopts an overall configuration that is drastically different from that observed in the Munc18-1-syntaxin-1 complex (Figs. 1D, 2D). Importantly, helices Hd and He of the linker form a small four-helix bundle with the N-terminus of the synaptobrevin SNARE motif and helix Hf of syntaxin-1 (Figs. 1E,F 2E,F) that is similar in class1 and class2 (Fig. 2J) and involves multiple contacts between synaptobrevin and the syntaxin-1 He helix (Fig. S6E-H). These observations suggest that the syntaxin-1 linker region acts as a template to nucleate SNARE complex assembly while Munc18-1 is a platform that controls SNARE zippering by holding the C-terminal halves of the syntaxin-1 and synaptobrevin SNARE motifs. This feature and the overall architectures observed in class1 and class2 explain why the syntaxin-1 N-terminal region is crucial for template complex formation (*30*).

### Alteration of Munc18-1-SNARE interactions by mutagenesis

To study the functional effects of disrupting the template complex and hence test the functional relevance of our cryo-EM structures, we used a battery of Munc18-1 mutations that included the gain-of-function D326K and P335A mutations. Note that the P335A mutation was previously designed to extend helix H12 and increase synaptobrevin binding, but such increase was not observed (*24*); instead, the P335A mutation decreased syntaxin-1 binding (*33*). To impair binding of Munc18-1 to syntaxin-1 at a different site, we used an S42Q mutation expected to disrupt interactions with the syntaxin-1 SNARE motif C-terminus. We also included the L307R and L348R mutations that disrupt synaptobrevin binding and/or template complex formation (*24, 30*); a E352K mutation that we designed to also disrupt Munc18-1-synaptobrevin binding; and a Q301D mutation that impairs neurotransmitter release and was reported to impair binding of Munc18-1 to synaptobrevin or to Munc13-1 (*34, 35*). The locations of the mutated residues are shown in Figs. 2A, S6A-D and S7A-C (D326 is visible in the Munc18-1-syntaxin complex but not in class1 and class2).

To analyze the effects of these mutations on binding of Munc18-1 to the syntaxin-1 cytoplasmic region and SyxLE/Syb, we used mass photometry, a single molecule interferometric scattering-based technique that allows counting of proteins and complexes existing in solution (*36*) (Fig. 3, Table S3). We measured a K_D_ of 3.3 nM between Munc18-1 and syntaxin-1, consistent with previous ITC data (*32*), whereas the K_D_ of Munc18-1 for SyxLE/Syb was 17.9 nM. The Munc18-1 P335A mutation induced Munc18-1 dimerization but we were able to distinguish the peak of the syntaxin-1-Munc18-1 P335A heterodimer (Fig. S8) and obtained a K_D_ of 12.3 nM, confirming that this mutation impairs syntaxin-1 binding (*33*). However, P335A had no overt effect on Munc18-1 binding to SyxLE/Syb (Fig. 3J). These results are consistent with the fact that P335 packs closely against syntaxin-1 in the Munc18-1-syntaxin-1 complex, whereas P335 does not contact syntaxin-1 in class1 and class2 (Fig. S6I-K). In contrast, the S42Q mutation markedly impaired binding of Munc18-1 to both syntaxin-1 and SyxLE/Syb (Fig. 3D,H,I,J), which indicates that the C-terminal half of the syntaxin-1 SNARE motif plays a key role in binding to Munc18-1 both in the binary complex and in the template complex, and is consistent with the similarity of the binding modes observed in the corresponding region of the binary complex, class1 and class2 (Fig. S7A-C). Binding of Munc18-1 to SyxLE/Syb was strengthened by the D326K mutation, as expected, and was impaired to different degrees by the L307R, Q301D, E352K and L348R mutations (Fig. 3J). These results are consistent with our cryo-EM structures and support the notion that the physiological effects of the Q301D mutation arise from impairment of Munc18-1 binding to synaptobrevin (*35*) rather than to Munc13-1 (*34*).

**Fig. 3.**
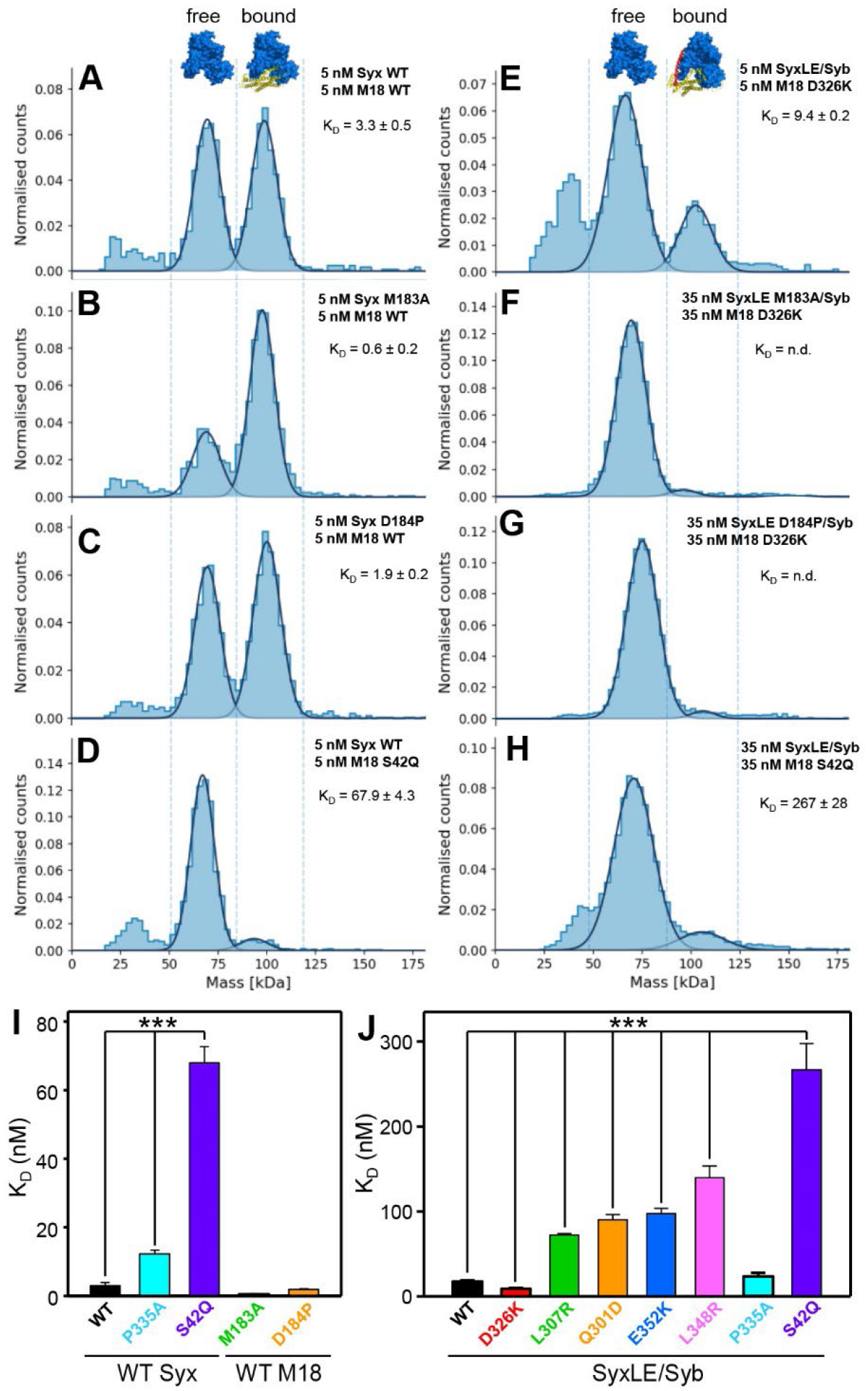
Analysis of Munc18-1-SNARE interactions by mass photometry. (**A**-**H**) Normalized histograms of mass distributions observed for samples containing the indicated concentrations of WT or mutant Munc18-1 (M18) plus WT or mutant syntaxin-1(2-253) (Syx) (**A**-**D**), or mutant Munc18-1 plus SyxLE/Syb with or without the M183A or D184P mutations (**E**-**H**). Gaussian fits (solid lines) were used to calculate the populations of free and bound Munc18-1, and derive dissociation constants (K_D_s). Binding of Munc18-1 D326K to SyxLE M183A/Syb and SyxLE D184P/Syb was too weak to derive reliable K_D_s. (**I**-**J**) Bar diagrams illustrating the average K_D_s (Table S3) obtained from six independent experiments for samples containing WT or mutant Munc18-1 plus WT or mutant syntaxin-1(2-253) (**I**), or WT or mutant Munc18-1 plus SyxLE/Syb with or without the M183A or D184P mutations (**J**). Error bars represent standard deviations. Statistical significance and p values were determined by one-way analysis of variance (ANOVA) with the Holm-Sidak test (*** p < 0.001).

### Disrupting the template complex impairs trans-SNARE complex formation and fusion

To test the effects of the Munc18-1 mutations on SNARE complex assembly, we first adapted a solution FRET assay which showed that the Munc13-1 MUN domain accelerates the transition from the Munc18-1-syntaxin-1 complex to the SNARE complex (*11*). To avoid using the high concentrations of MUN domain required for such acceleration, we covalently attached the MUN domain to the C-terminus of the syntaxin-1 cytoplasmic region with a long flexible linker (Fig. 4A). The syntaxin-1-MUN domain fusion (SyxMUN) assembled into SNARE complexes with SNAP-25 and soluble synaptobrevin as efficiently as syntaxin-1 alone (Fig. 4B). Binding of Munc18-1 to syntaxin-1 alone abolished SNARE complex assembly, but the SyxMUN fusion bound to Munc18-1 was still able to assemble into SNARE complexes (Fig. 4B). The reaction was substantially stimulated by the P335A and S42Q Munc18-1 mutations but not by D326K, and slight accelerations also appeared to be caused by the mutations that disrupt synaptobrevin binding (Figs. 4C, S9A). The P335A and S42Q mutations were able to allow SNARE complex assembly even in the absence of the MUN domain (Figs. 4D, S9B). These results indicate that the rate-limiting step in SNARE complex assembly in these solution assays is not the binding of synaptobrevin to the Munc18-1-syntaxin-1 complex to form the template complex but rather the dissociation of the syntaxin-1 SNARE motif from Munc18-1.

**Fig. 4.**
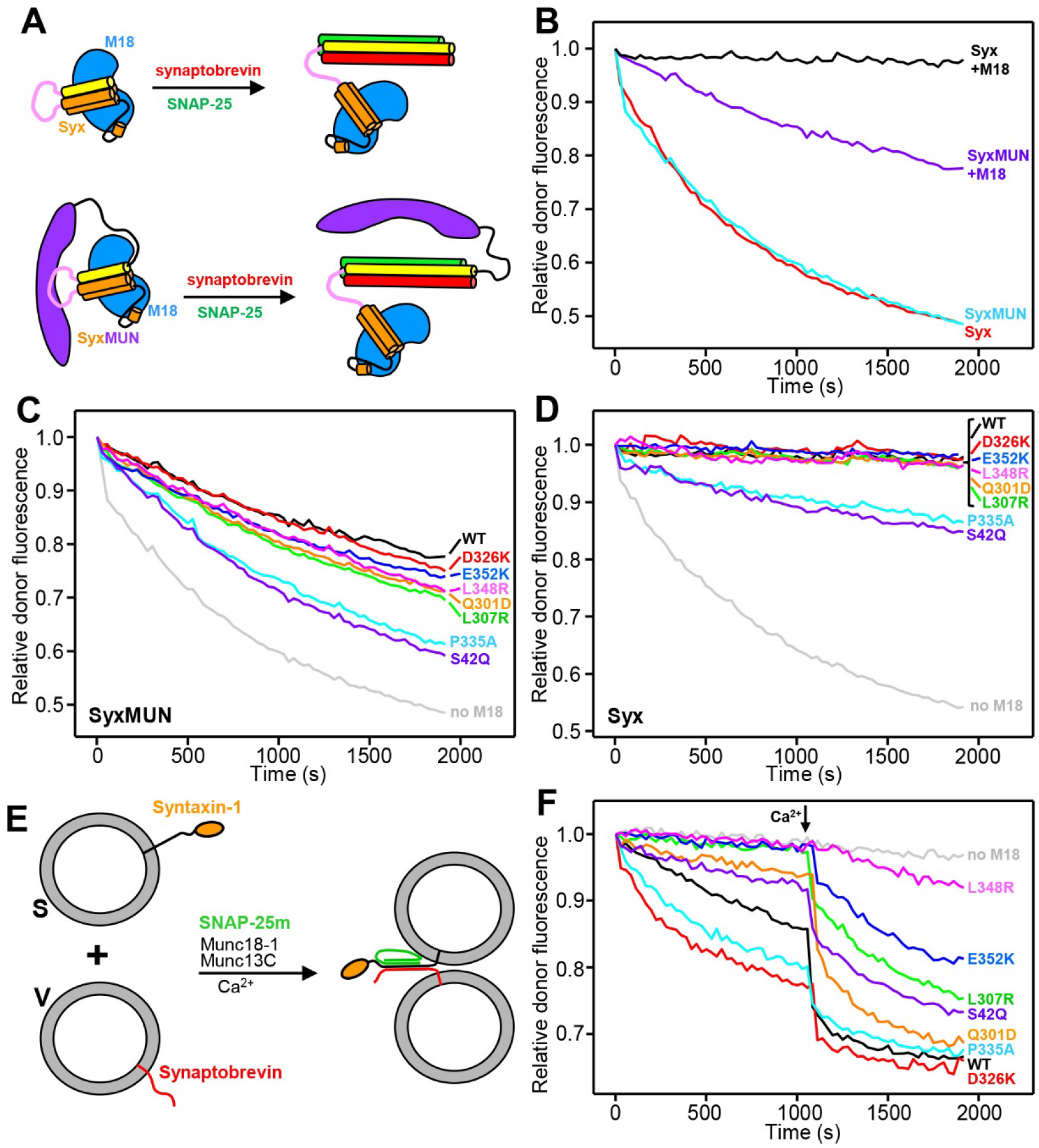
Effects of mutations in Munc18-1 on SNARE complex assembly rates. (**A**) Diagram summarizing the assays used to monitor SNARE complex assembly in solution starting with Munc18-1 (blue) bound to the syntaxin-1 cytoplasmic region (H_abc_ domain orange, linker pink, SNARE motif yellow) alone (Syx) or covalently linked to the Munc13-1 MUN domain (purple) (SyxMUN). Synaptobrevin (red) was labeled with a donor fluorescent probe and SNAP-25 (green) was labeled with an acceptor probe. SNARE complex assembly was monitored by the development of FRET between the probes. (**B**-**D**) Decrease in relative donor fluorescence (normalized to the first time point) as a function of time in SNARE complex assembly reactions where synaptobrevin and SNAP-25 were added to Syx or SyxMUN free or pre-bound to WT Munc18-1 (M18) (**B**), to SyxMUN bound to WT or mutant Munc18-1 (**C**) or to Syx bound to WT or mutant Munc18-1 (as indicated) (**D**). (**E**) Diagram summarizing trans-SNARE complex assembly assays between liposomes containing synaptobrevin labeled with a donor fluorescent probe (V-liposomes) and liposomes containing syntaxin-1 labeled with an acceptor probe (S-liposomes) in the presence of Munc13C, a SNAP-25 mutant (SNAP-25m) and WT or mutant Munc18-1. (**F**) Decrease in relative donor fluorescence (normalized to the first time point) as a function of time in trans-SNARE complex assembly assays performed with WT or mutant Munc18-1 as indicated. Reactions were initiated in the presence of 100 μM EGTA and Ca^2+^ (600 μM) was added at the time indicated by the arrow.

To better mimic the geometry of SNARE complex assembly in neurons, we next used a FRET assay that measures assembly of trans-SNARE complexes between synaptobrevin-containing liposomes (V-liposomes) and syntaxin-1 liposomes (S-liposomes) (Fig. 4E), adapting a previously described assay that used SNAP-25 with a mutation in the C-terminus of its first SNARE motif (SNAP-25m) to prevent liposome fusion (*10*). The S-liposomes were incubated with WT or mutant Munc18-1, and trans-SNARE complex assembly was monitored upon addition of V-liposomes, SNAP-25m and a fragment spanning the conserved C-terminal region of Munc13-1 (Munc13C). As expected (*10*), assembly in the presence of WT Munc18-1 was slow in the absence of Ca^2+^ and was strongly activated by Ca^2+^, which likely arises because Ca^2+^ binding to the Munc13-1 C_2_B domain changes the orientation in which Munc13C bridges the two membranes (*37*). The D326K and P335A mutations substantially enhanced Ca^2+^-independent trans-SNARE complex assembly (Figs. 4F, S9C), consistent with the gains-of-function observed previously for these two mutants (*16, 29*). However, the S42Q mutation impaired trans-SNARE complex assembly, in contrast to the effect observed for this mutation in the solution assay. The Q301D, L307R, L348R and E352K mutations decreased assembly in the absence and presence of Ca^2+^ to different extents (Figs. 4F, S9C,D) that approximately correlated with impairment in binding of Munc18-1 to SyxLE/Syb (Fig. 3J).

To analyze how the mutations affect the ability of Munc18-1 to support SNARE-dependent membrane fusion, we used an assay that measures content mixing between V-liposomes and liposomes containing syntaxin-1 and SNAP-25 (T-liposomes) in the presence of Munc18-1, Munc13C, NSF and αSNAP (*13*), and that has allowed us to establish many correlations between the effects of mutations on liposome fusion and their effects on neurotransmitter release (*13, 14, 37, 38*) (Fig. 5A). As expected, no liposome fusion was observed in the absence of Munc18-1 and inclusion of WT Munc18-1 yielded inefficient Ca^2+^-independent fusion that was dramatically stimulated by addition of Ca^2+^ (Fig. 5B). The D326K and P335A mutations strongly enhanced Ca^2+^-independent fusion, as observed previously (*16, 29*), and the S42Q induced a similar stimulation (Figs. 5B, S10A). In contrast, the Q301D, L307R, L348R and E352K mutations abolished the small amount of Ca^2+^-independent fusion observed for WT Munc18-1 and impaired Ca^2+^- dependent fusion to different extents (Figs. 5B, S10A,B) that approximately correlate with the effects of these mutations on binding of Munc18-1 to SyxLE/Syb and on trans-SNARE complex assembly (Figs. 3J, 4F, S9C,D). Although the effects of the L307R and Q301D mutations on Ca^2+^-dependent fusion appear to be small, they are significant, as mild impairments in activity in these assays are often not observable (*37*).

**Fig. 5.**
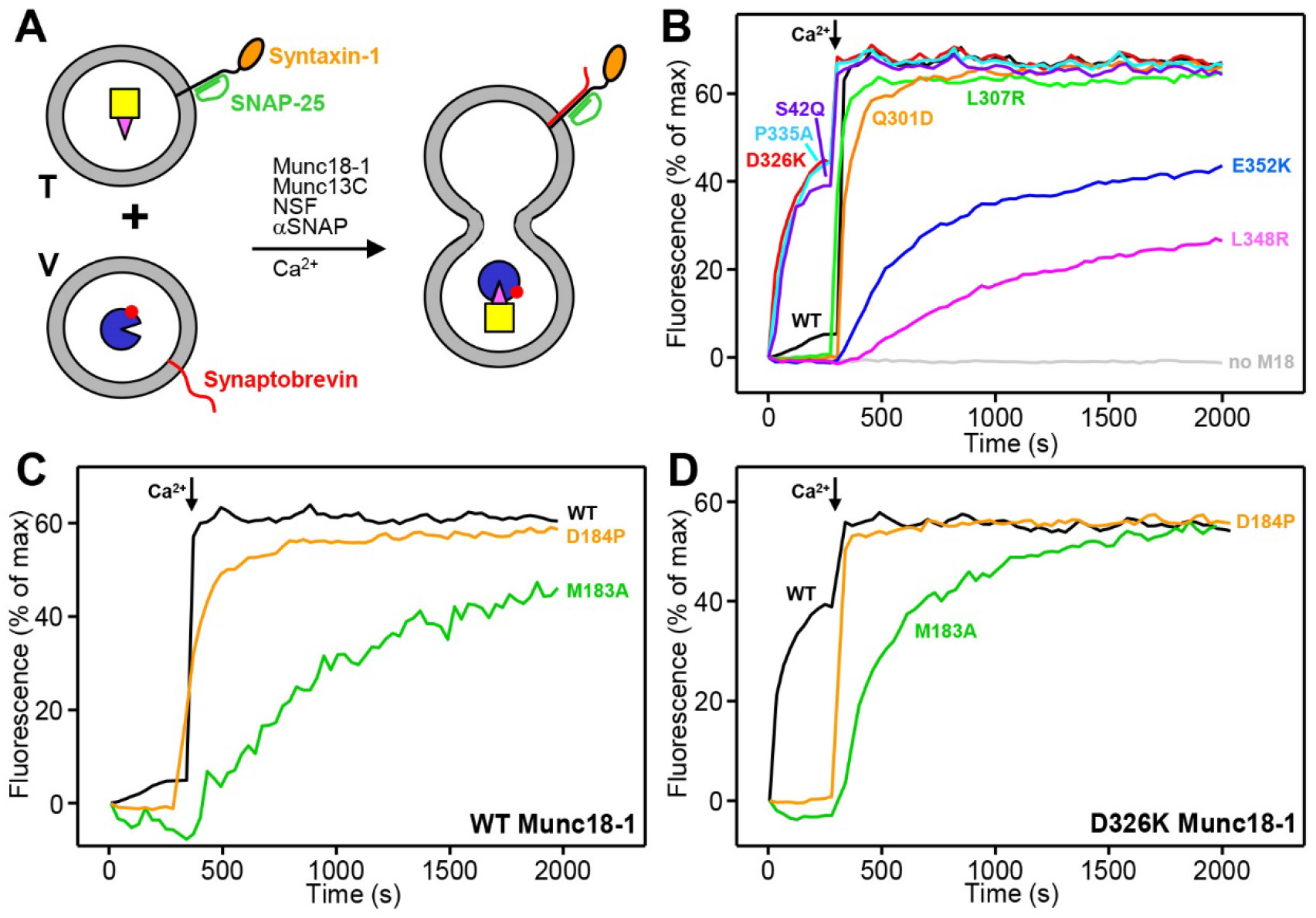
Effects of Munc18-1 and syntaxin-1 mutations on liposome fusion. (**A**) Diagram summarizing the content mixing between T-liposomes containing syntaxin-1 and SNAP-25, which were pre-incubated with Munc18-1, NSF and αSNAP, and V-liposomes containing synaptobrevin in the presence of Munc13C. V-liposomes contain trapped Cy5-strepatavidin and T-liposomes contain trapped PhycoE-biotin. (**B-D**) Content mixing between V-liposomes and T-liposomes was monitored from the increase in the fluorescence signal of Cy5-streptavidin caused by FRET with PhycoE-biotin. Assays were performed with WT or mutant Munc18-1 (**B**), or WT or mutant syntaxin-1 and WT (**C**) or D326K mutant (**D**) Munc18-1, as indicated. Experiments were started in the presence of 100 μM EGTA, and Ca2+ (600 μM) was added at 300 s.

The formation of a small four-helix bundle by synaptobrevin, the syntaxin-1 SNARE motif and helices Hd and He from the syntaxin-1 linker in class1 and class2 (Figs. 1E,F 2E,F) suggests that the linker acts as a template to initiate assembly of the SNARE complex. To test the functional relevance of the structure formed in this region, we designed two mutations in helix He: a D184P mutation expected to disrupt the helical structure and an M183A mutation designed to disrupt hydrophobic contacts that stabilize the short four-helix bundle (Fig. S6E-H) and involve a highly conserved residue (Fig. S11). Importantly, these mutations are not expected to substantially perturb the closed conformation of syntaxin-1 because these residues are exposed in a loop (Fig. S7G). Indeed, the M183A and D184P mutations did not impair syntaxin-1 binding to Munc18-1 but, interestingly, they strongly disrupted binding of SyxLE/Syb to Munc18-1 D326K (Fig. 3A-C,E-G). Moreover, Ca^2+^-dependent liposome fusion in the presence of WT Munc18-1 was strongly disrupted by the M183A mutation and was impaired by the D184P mutation (Figs. 5C, S10C). Analysis of the liposomes by SDS-PAGE (Fig. S10E) showed that these results did not arise from poor syntaxin-1 incorporation onto the liposomes. Since the high efficiency of Ca^2+^-dependent fusion might have masked the severity of the effect of the D184P mutation, we also performed liposome fusion assays in the presence of the Munc18-1 D326K mutant, which yields Ca^2+^-independent fusion. Both M183A and D184P abolished Ca^2+^-independent fusion, emphasizing the importance of the He helix of syntaxin-1 for the molecular events that lead to liposome fusion.

## Discussion

The crystals structures of Vps33-SNARE complexes (*28*) and multiple data on Munc18-1 (*24, 29, 30*) showed that SM proteins form templates for SNARE complex assembly and that a synaptobrevin-Munc18-1-syntaxin-1 template complex represents a crucial intermediate in the path from the Munc18-1-closed syntaxin-1 complex to the SNARE complex. However, the steps leading to SNARE complex assembly and the basis for the key role played by the syntaxin-1 N-terminal region in template complex formation remained poorly understood, and no structure of an SM protein bound to its two cognate SNAREs was available. Our cryo-EM structures now fill this gap and, together with our biophysical assays and previous data, allow us to develop a realistic model of the steps that lead to the SNARE complex in which the syntaxin-1 linker plays a key role in nucleating SNARE assembly while the C-termini of the syntaxin-1 and synaptobrevin SNARE motifs are held at distant sites of Munc18-1.

Structural characterization of the template complex was hindered by its necessarily dynamic nature as an intermediate in the pathway to SNARE complex assembly. To overcome this problem, we used chemical cross-linking and took advantage of the ability of cryo-EM to characterize substantially populated states within conformational ensembles. In principle, the introduction of a disulfide bond between synaptobrevin and syntaxin-1 might cause local structural distortions, but the bias introduced by the cross-link most likely selects for configurations of the ensemble that are on the productive pathway for SNARE complex assembly. First, the two cross-linked residues are in contact in the structure of the SNARE complex (*3*). Second, the cross-link facilitated SNARE complex assembly in our SDS-PAGE assay (Fig. S1C) and formation of the template complex in optical tweezer experiments (*30*). Third, these experiments established multiple correlations between the effects of mutations on cross-linked template complex formation and their effects on liposome fusion or neurotransmitter release (*30*). And fourth, our cryo-EM structures readily explain our Munc18-1-SNARE binding, SNARE complex assembly and liposome fusion data. Thus, the effects of Munc18-1 mutations on binding (Fig. 3) support the interaction modes of Munc18-1 with synaptobrevin and syntaxin-1 observed in the two structures. Moreover, the effects of the Munc18-1 mutations on SNARE complex assembly and fusion (Figs. 4-5) illustrate the importance not only of Munc18-1 binding to the syntaxin-1 and synaptobrevin SNARE motifs but also of release of these interactions to allow SNARE complex formation (supplementary discussion). Importantly, compelling evidence for the functional relevance of the conformations observed for the syntaxin-1 linker in our cryo-

EM structures is provided by the severe disruption of Munc18-1-SyxLE/Syb binding and of liposome fusion caused by the M183A and D184P mutations in syntaxin-1 (Figs. 3, 5C,D).

Our results suggest a model for the transition from the Munc18-1-syntaxin-1 complex to the SNARE complex where states resembling our two cryo-EM structures of the template complex are at center stage (Fig. 6). The interactions of Munc18-1 with the N-peptide and the C-terminal half of the SNARE motif of syntaxin-1 in the Munc18-1-syntaxin-1 complex, which are also present in class1 and class2 (Figs. 1D-F, S4A-C), provide anchor points to keep syntaxin-1 bound to Munc18-1 while the N-terminal half of the SNARE motif and the linker undergo conformational rearrangements that are necessary to form the template complex. Unfurling of the Munc18-1 loop allows binding of the C-terminal half of the synaptobrevin SNARE motif (Figs. 2A-C, 6B), yielding an anchor point for synaptobrevin. Thus, a central aspect of this model is that the attachment of the C-terminal halves of the synaptobrevin and syntaxin-1 SNARE motifs to distal parts of Munc18-1 allows their N-terminal halves to come close and bind, as proposed from optical tweezer experiments (*30*). Such binding requires release of interactions of the syntaxin-1 SNARE motif with the H_abc_ domain and with a region of Munc18-1 containing P335 and a β-hairpin (Figs. 2A-B, S6I-K, S7H-J), which imposes an energy barrier that may be difficult to overcome with just a few synaptobrevin-syntaxin-1 interactions. Remodeling of the syntaxin-1 linker to form helices Hd and He, and packing of these helices against the syntaxin-1 and synaptobrevin SNARE motifs (Figs. 2E,F, S6E-H), likely helps to overcome this energy barrier.

**Fig. 6.**
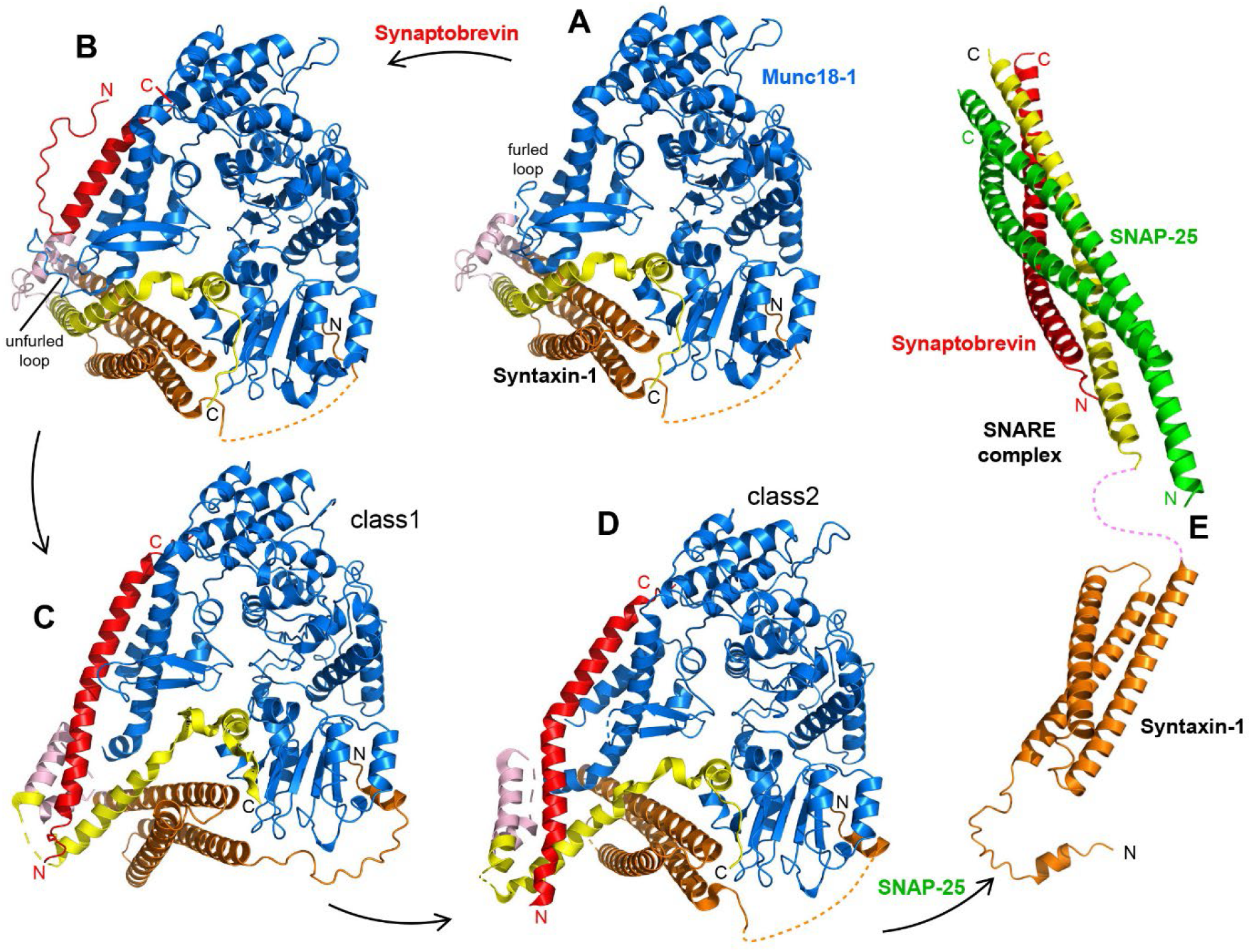
Model of SNARE complex assembly templated by Munc18-1 and the syntaxin-1 N-terminal region. The model postulates that assembly starts with Munc18-1 bound to closed syntaxin-1 (**A**) (PDB code 3C98) and is initiated when the Munc18-1 loop unfurls to allow synaptobrevin binding (**B**) (model built manually). Conformational changes in syntaxin-1 that are stimulated by Munc13-1 (not shown), leads to template complex configurations such as those of class1 and class2, where syntaxin-1 gradually opens and the syntaxin-1 linker nucleates interactions between the syntaxin-1 and synaptobrevin SNARE motifs (**C**,**D**). The color code is the same as in Fig. 1. Binding to SNAP-25 (green) eventually leads to formation of the SNARE complex (**E**) (PDB codes: H_abc_ domain 1BR0; SNARE four-helix bundle 1SFC). The N-and C-termini of the SNAREs are indicated in the relevant panels.

The resulting small four-helix bundle includes native contacts between residues of synaptobrevin and syntaxin-1 (e.g. V42, M46 of synaptobrevin and L212, F216 of syntaxin-1; Fig. S6E,F) that also interact in the SNARE complex and is therefore likely to constitute the initiation point for SNARE complex formation. The different extents of interaction between the syntaxin-1 SNARE motif and the H_abc_ domain observed in the Munc18-1-syntaxin-1 complex, class1 and class2 (Figs. 2D-F, 6A,C,D, S7H-J) help to visualize how syntaxin-1 opens gradually and suggest that, once the small four-helix bundles forms, it helps to maintain the native synaptobrevin-syntaxin-1 contacts during the opening process. However, this structure is likely metastable and helices Hd-He need to be at least partially dissociated in subsequent events to allow SNAP-25 binding to syntaxin-1 and synaptobrevin. As the SNARE complex zippers, the C-terminal halves of the synaptobrevin and syntaxin-1 SNARE motifs need to dissociate from Munc18-1 to allow full SNARE complex assembly (Fig. 6E). Hence, in this model, the syntaxin-1 linker plays a central role in templating SNARE complex assembly together with Munc18-1 in this model.

Some of these events may occur in different order (supplementary discussion) but, regardless of the order, it is clear from our results and previous data that there are multiple energy barriers that hinder the transition from the Munc18-1-syntaxin-1 complex to the template complex and then to the SNARE complex, providing varied possibilities for regulation. Indeed, Munc13-1 acts as a master regulator of release (*5*) at least in part because it controls SNARE complex assembly by bridging the two membranes in more than one orientation (*14, 37*) and helping to open syntaxin-1 (*11, 12*) through weak interactions with the linker (*39, 40*) that are expected to influence the molecular events outlined above. Release is also modulated by kinases that phosphorylate Munc18-1, altering interactions with SNAREs [e.g. (*41*)]. Hence, this pathway is central for the exquisite regulation of synaptic vesicle fusion, which is a hallmark that allows presynaptic terminals to act as small computational units in the brain (*42*). The facts that a mutation in Unc-18 analogous to P335A partially rescues the dramatic phenotypes caused by deletion of Unc-13 (*16*) and that the LE mutation of syntaxin-1 can partially rescue the phenotypes caused by absence of very diverse proteins (*15, 22, 23*) emphasize how understanding the mechanism of neurotransmitter release allows manipulation of the release efficiency. The structures of Munc18-1-SNARE complexes that are now available, together with the mechanistic understanding yielded by our mutagenesis studies (Figs. 3-5) and previous data [reviewed in (*5*)], provide a framework to design strategies for modulation of neurotransmitter release and other forms of regulated secretion that might have therapeutic use for a wide variety of diseases.

## Author contributions

KPS, XB and JR conceived the research; KPS conducted all experiments; KPS and JR analyzed the data; JX froze the cryo-EM grids; XB and JX acquired the cryo-EM data; XB processed and analyzed the cryo-EM data; KPS and XZ built and refined the structural models; JR and KPS wrote the draft, with input from all authors.

## Data and materials availability

All data are available in the main text or the supplementary materials. The two cryo-EM structures are been deposited in the Protein Data Bank.

## Supplementary materials

Materials and Methods Supplementary Text

Figs. S1 to S11

Tables S1 to S3

References 1-51

