## Supplementary materials for "SNARE assembly enlightened by cryo-EM structures of a synaptobrevin-Munc18-1-syntaxin-1 complex"

**Pdf contains**

Materials and methods

Supplementary text

Figs. S1-S11

Tables S1-S3

References 1-51

### Materials and methods

**Protein expression and purification.** *E. coli* expression and purification of full-length rat syntaxin-1A, the cytoplasmic fragment of rat syntaxin-1A (residues 2-253), a cysteine-free variant of full-length rat SNAP-25a, a full-length rat synaptobrevin-2, rat synaptobrevin-2 (residues 29-83), rat synaptobrevin (residues 1-96), full-length rat Munc18-1, full length *Cricetulus griseus* NSF, full-length *Bos Taurus* αSNAP, and a rat Munc13-1 fragment spanning the $C_1$, $C_2B$, MUN and $C_2C$ domains (residues 529-1725, Δ1408-1452) (referred to as Munc13C) were described previously (*9, 10, 13, 43-45*). The following mutants were also described previously (*10, 29, 40*) and were purified through the same protocols used for the WT proteins: full-length syntaxin-1 S186C, C145A, C271A, C272A; SNAP-25a R136C C84S, C85S, C90S, C92S; SNAP-25a M71D, L78D C84S, C85S, C90S, C92S; full-length synaptobrevin-2 L26C; synaptobrevin-1 1-96 L26C; Munc18-1 P335A; Munc18-1 D326K; Munc18-1 L348R.

Briefly, full-length syntaxin-1 was expressed overnight at 25 °C upon induction with 0.4 mM isopropyl-β-D-thiogalactoside (IPTG). Cell pellets were re-suspended with 20 mM 4-(2-hydroxyethyl)-1-piperazineethanesulfonic acid (HEPES), pH 7.4, 500 mM NaCl, 8 mM imidazole, 1 mM tris(2-carboxyethyl)phosphine (TCEP). The protein was initially purified via affinity chromatography using HisPur Ni-NTA resin (Thermo Fisher) in 20 mM tris(hydroxymethyl)aminomethane (Tris), pH 7.4, 500 mM NaCl, 8 mM imidazole, 2% Triton X-100 (v/v), 6 M urea. Upon extensive washes, the protein was eluted in 20 mM Tris, pH 7.4, 500 mM NaCl, 400 mM imidazole, and 0.1% dodecylphosphocholine (DPC). The polyhistidine tag was removed using thrombin protease, followed by size-exclusion chromatography on a Superdex 200 column (GE 10/300) equilibrated in 20 mM Tris, pH 7.4, 125 mM NaCl, 1 mM TCEP, and 0.2% DPC.

Expression of syntaxin-1 2–253 was induced with 0.4 mM IPTG and expressed overnight at 25 °C. Upon cell lysis, purification was done using Glutathione Sepharose 4B resin (GE) in phosphate-buffer saline pH 7.4 (PBS), PBS with 1% Triton X-100 (v/v), and PBS with 1 M NaCl. The GST-tag was cleaved by thrombin protease and the eluted protein was further purified by anion exchange chromatography on a HiTrap Q column (GE) in 25 mM Tris, pH 7.4, and 1 mM TCEP using a linear gradient from 0 to 1 M NaCl. Please note that the purification of syntaxin-1 2-253 C145A, L165A, E166A, L205C (SyxLE) was carried with the same procedure except that the final purification was done by size exclusion chromatography using a Superdex 75 (GE 10/300) column equilibrated withs 100 mM phosphate buffer at pH 7.0 with 100 mM NaCl, 6 M urea, 0.2 mM ethylenediaminetetraacetic acid (EDTA) and 0.5 mM TCEP.

Cysteine-free SNAP-25 was expressed overnight at 25 °C upon induction with 0.4 mM IPTG. Upon cell lysis, protein purification was performed using HisPur Ni-NTA resin (Thermo Fisher) in 50 mM Tris, pH 8.0, 500 mM NaCl, 20 mM imidazole, and 1% Triton X-100 (v/v). The His6-tag was cleaved by thrombin protease, and the protein was purified by size-exclusion chromatography using a Superdex 75 column (GE 16/60) in 50 mM Tris, pH 8.0, and 150 mM NaCl.

Full-length synaptobrevin-2 was expressed overnight at 25 °C upon induction with 0.4 mM IPTG. Cells were re-suspended in PBS buffer containing 1% Triton X-100 (v/v). Purification was done using Glutathione Sepharose 4B resin (GE) at 4 °C. The bound proteins were treated with PBS with 1% Triton X-100 (v/v), followed by the addition of thrombin to cleave the GST-tag. The protein was further purified by cation exchange chromatography on a HiTrap S column (GE) in 25 mM NaAc, pH 5.5, 1 mM TCEP, and 1% (w/v) using a linear gradient from 0 to 1 M NaCl.

Expression of synaptobrevin-2 1-96 or synaptobrevin-2 29-83 Q36C (Syb) was induced with 0.4 mM IPTG and expressed overnight at 23 °C. Purification was done using Glutathione Sepharose 4B resin (GE), followed by cleavage of the GST-tag. The final purification of synaptobrevin-2 1-96 was performed using size-exclusion chromatography on a Superdex 75 column (GE 16/60) equilibrated in 20 mM Tris, pH 7.4, and 125 mM NaCl. The final purification of synaptobrevin-2 29-83 Q36C was carried out using size-exclusion chromatography on a Superdex 75 column (GE 10/300) equilibrated with 100 mM phosphate buffer at pH 7.0 with 100 mM NaCl, 6M urea, 0.2 mM EDTA and 0.5 mM TCEP.

Expression of αSNAP was induced by the addition of 0.4 mM ITPG and continued overnight at 25 °C. Protein purification was performed using Glutathione Sepharose 4B resin (GE) by washing bound proteins with PBS, PBS with 1% Triton X-100 (v/v), and PBS with 1 M NaCl. Upon GST-tag cleavage in the presence of thrombin, the protein was purified by size-exclusion chromatography using a Superdex 75 column (GE 16/60) in 20 mM Tris, pH 7.4, 150 mM KCl, and 1 mM TCEP.

Expression of NSF was induced with 0.4 mM IPTG and continued overnight at 20 °C. Purification was performed using HisPur Ni-NTA resin (Thermo Fisher), followed by size-exclusion chromatography of hexameric NSF on a Superdex S200 column (GE 16/60) in 50 mM Tris, pH 8.0, 100 mM NaCl, 1 mM adenosine triphosphate (ATP), 1 mM EDTA, 1 mM dithiothreitol (DTT), and 10% glycerol (v/v). Removal of the His6-tag and monomerization of NSF was performed using TEV protease and apyrase, respectively, while dialyzing with nucleotide-free buffer for 36 h. To separate the hexameric form of NSF from the monomeric, three rounds of size-exclusion chromatography on a Superdex S200 column (GE 16/60) in 50 mM NaPi, pH 8.0, 100 mM NaCl, and 0.5 mM TCEP were performed by re-injecting fractions with hexameric NSF. Final reassembly of monomers and gel filtration chromatography of reassembled hexameric NSF were done using a Superdex S200 column (GE 16/60) in 50 mM Tris, pH 8.0, 100 mM NaCl, 1 mM ATP, 1 mM EDTA, 1 mM TCEP, and 10% glycerol (v/v).

Expression of full-length Munc18-1 was inducted with 0.4 mM IPTG and continued overnight at 20 °C. Upon cell lysis and centrifugation, the supernatant was loaded on Glutathione Sepharose 4B resin (GE) at 4 °C and the bound proteins were washed with PBS, PBS with 1% Triton X-100 (v/v), and PBS with 1 M NaCl. The GST-tag was cleaved by a thrombin treatment, followed by immediate size-exclusion chromatography using a Superdex 200 column (GE 16/60) in a buffer containing 20 mM Tris, pH 7.4, 200 mM KCl, and 1 mM TCEP.

Expression of the rat Munc13C (residues 529–1725, Δ1408–1452) was induced with 0.5 mM IPTG and performed overnight at 16 °C. The protein was purified by affinity chromatography using HisPur Ni-

NTA resin (Thermo Fisher) with extensive washes with 50 mM Tris, pH 8, 10 mM imidazole, 750 mM NaCl, 1 mM TCEP, and 10% glycerol (v/v). The protein was eluted with 50 mM Tris, pH 8, 250 mM NaCl, 1 mM TCEP, 10% glycerol (v/v), 500 mM imidazole and dialyzed overnight at 4 °C in 50 mM Tris, pH 8, 250 mM NaCl, 1 mM TCEP, 2.5 mM $CaCl_2$, 10% glycerol (v/v) in the presence of thrombin. The protein was further purified by anion exchange chromatography on a HiTrap Q column (GE) in 20 mM Tris, pH 8.0, 1 mM TCEP, and 10% glycerol (v/v) using a linear gradient from 0 to 1 M NaCl.

The following mutants were generated using the QuickChange site-directed mutagenesis and custom-designed primers and purified as the unmodified constructs: full-length syntaxin-1A (1-288) M138A, full-length syntaxin-1A (1-288) D184P, syntaxin-1A (2-253) M183A, syntaxin-1A (2-253) D184P, syntaxin-1A (2-253) C145A, L165A, E166A, L205C, syntaxin-1A (2-253) C145A, L165A, E166A M183A L205C, full-length syntaxin-1A (1-288) C145A, L165A, E166A, M183A, L205C, full-length syntaxin-1A (1-288) C145A, L165A, E166A, D184P, L205C, synaptobrevin-2 29-83 Q36C, Munc18-1 S42Q, Munc18-1 L307R, Munc18-1 Q301D and Munc18-1 E352K.

**Template complex formation.** Prior to complex formation, TCEP was removed from each protein preparation using Superdex 75 10/300 GL column. TCEP-free Syb (100-200 µM) was mixed with 400-500 µM of 2,2-dithiodipyridine (Sigma). The reaction was monitored at 343 nm using a Agilent 8453 UV-visible spectrometer. Upon completion, the excess of 2,2-dithiodipyridine was removed using a PD miniTrap G-25 Sephadex column with 100 mM phosphate at pH 7.0, 100 mM NaCl, 6 M urea, 0.2 mM EDTA as the elution buffer. An excess of activated Syb was then mixed with TCEP-free SyxLE and left incubated at room temperature overnight. The efficiency of disulfide bond formation between Syb and SyxLE was estimated based on the absorbance at 343 nm. The SyxLE/Syb conjugate was purified by size-exclusion chromatography using Superdex 75 10/300 GL and 20 mM HEPES of pH 7.4, 150 mM KCl as the running buffer. TCEP-free Munc18-1 D326K was then mixed with purified SyxLE/Syb and left rotated overnight at room temperature. The template complex was finally purified by size-exclusion chromatography using a Superdex 75 10/300 GL column in 20 mM HEPES of pH 7.4 with 150 mM KCl.

**EM data acquisition**

Flourinated fos-choline-8 was added into purified template complex formed by SyxLE and Munc18-1 D326K (7.5 mg/ml) to a final concentration of 5 mM and 3 µl of the sample were applied to glow-discharged (30 mA, 80 s) Quantifoil R1.2/1.3 300-mesh gold holey carbon grids (Quantifoil, Micro Tools GmbH, Germany). Grids were blotted for 4.0 seconds under 100% humidity at 4 °C before being plunged into the liquid ethane using a Mark IV Vitrobot (FEI). Micrographs were acquired on a Titan Krios microscope (FEI) operated at 300 kV with a K3 direct electron detector (Gatan), using a slit width of 20 eV on a GIF-Quantum energy filter. SerialEM was used for data collection. A calibrated magnification of 46,296 was used for imaging of the samples, yielding a pixel size of 1.08 Å on the images. The defocus range was set from -1.6 µm to -2.6 µm. Each micrograph was dose-fractionated to 30 frames with a total dose of about 60 e-/$Å^2$.

**Image processing**

The cryo-EM refinement statistics are summarized in Table S1. 7,401 movie frames of the SyxLE/Syb-Munc18-1 D326K complex were motion-corrected and binned two-fold, resulting in a pixel size of 1.08 Å, and dose-weighted using MotionCor2. The CTF parameters were estimated using Gctf. RELION3 (*46*) was used for the following processing. Particles were first roughly picked by using the Laplacian-of-Gaussian blob method, and then subjected to 2D classification. Class averages representing projections of the SyxLE/Syb-Munc18-1 D326 complex in different orientations were used as templates for reference-based particle picking. Extracted particles were binned three times and subjected to 2D classification. Particles from the classes with fine structural features were selected for 3D classification using an initial model generated from a subset of the particles in RELION3. Particles from one of the resulting 3D classes showing good secondary structural features were selected and re-extracted into the original pixel size of 1.08 Å. Subsequently, we performed finer 3D classification imposed by using local search in combination with small angular sampling (3.75º), resulting in new classes showing two distinct conformations of the SyxLE/Syb-Munc18-1 D326 complex. The particles corresponding to two different conformations were selected and refined separately, leading to two different cryo-EM maps at 3.5 Å (class2) and 3.7 Å (class1) resolution, respectively.

**Model building and refinement.** Model building of both template complexes was started by docking the individual chains from the previously solved crystal structure of the syntaxin-1-Munc18-1 complex (PDB ID: 3C98) using Chimera 1.15 (*47*). The models were improved by iterative manual building in Coot 0.9 (*48*) and real-space refinement in the software package Phenix 1.19.1 (*49*). The density for He in syntaxin in both tc1 and tc2 is rather weak, preventing the assignment of individual residues. This part was therefore built as a poly-alanine helix, with the register of the residues undetermined. The quality of the model stereochemistry and fit to density was evaluated with the comprehensive validation method Molprobity (*50*) as implemented in the Phenix package. The statistics are shown in Table S1.

**Gel filtration binding assay.** 6-20 μM samples containing Munc18-1 D326K, or Munc18-1 D326K incubated with SyxLE and Syb, or Munc18-1 D326K incubated with SyxLE/Syb were injected into a size exclusion chromatography column Superdex 30 Increase 10/300 GL using 20 mM HEPES buffer, pH 7.4, 150 mM KCl as running buffer. To form potential complexes selected proteins were incubated overnight at 4 ˚C before injection. Selected eluted fractions were loaded into SDS-PAGE gel and stained with InstantBlue Coomassie protein stain to confirm protein co-elution.

**SDS-PAGE SNARE complex assembly assay.** To detect the SDS-resistant SNARE complex, the template complex was formed as described above and the SyxLE/Munc18-1 D326K complex was preformed (overnight incubation at 4 °C). 5 µM SNAP-25 was added to 5 µM template complex, or 5 µM Syb and 5 µM SNAP-25 were added to the SyxLE/Munc18-1 D326K complex, or 5 µM Syb, 5 µM SNAP-25 and 5 µM SyxLE were mixed. The reaction was done in 25 mM HEPES pH 7.4, 150 mM KCl, 10% glycerol (v/v) at room temperature. After 3 minutes, the reaction was stopped by addition of the SDS-PAGE gel-loading buffer. The samples were then loaded into SDS-PAGE gels and stained with InstantBlue Coomassie protein stain. The gels were imaged using a Bio-Rad ChemiDoc imaging system.

**Mass photometry.** Samples containing 1000 nM SyxLE/Syb, or SyxLE M183A/Syb, or SyxLE D184P/Syb, or syntaxin-1 (2-253), or syntaxin-1 (2-253) M183A, syntaxin-1 (2-253) D184P were incubated overnight at room temperate with TCEP-free 1000 nM or 2000 nM WT or mutant Munc18-1. Prior to the measurements the samples were diluted tenfold using PBS at pH 7.4. High precision microscope cover glasses were rinsed with Mili-Q water, isopropanol, Mili-Q water, isopropanol, and Mili-Q water, and dried using a stream of nitrogen gas. Clean coverslips with attached silicon gaskets were then mounted on immersion oil (refractive index of 1.518)-covered lenses of a Refeyn's second-generation mass photometer. All the measurements were performed using PBS at pH 7.4. Data were collected using AcquireMP software. A single gasket was filled with 13-17.1 µL of phosphate-buffered saline to enable focusing of the coverslip. Once the focus signal was stable, 0.9-7 µL of the 100 nM sample was added and quickly mixed to achieve the desired final protein concentrations. A movie was recorded for 60 s (2819 frames) and processed using DiscoverMP. Contrast-to-mass calibration was achieved using a 0.002 mg/ml BSA standard. The contrasts observed for BSA monomer (66 kDa), dimer (132 kDa) and trimer (198 kDa) were used to generate a standard calibration curve. Each measurement is displayed as normalized histograms with Gaussian fitting of the binding event counts with a loaded standard calibration curve. The $K_D$ was calculated using a standard one-ligand binding model.

**Solution SNARE complex assembly assay.** To monitor SNARE complex assembly in solution, synaptobrevin (1-96) L26C and SNAP-25 R136C (50-150 μM) were respectively labeled with Alexa-488 maleimide and tetramethylrhodamine (TMR) maleimide at room temperature with 10-20 fold excess of the dyes for 2 hours. The excess of the reagents was removed using a Superdex 75 (10/300) column equilibrated with 20 mM Tris pH 7.4, 150 mM NaCl, 1 mM TCEP. SNARE complex assembly was monitored by detecting Alexa-488 donor fluorescence intensity (excitation at 468 nm, emission at 518 nm) as a function of time at 37 °C using a PTI Quantamaster 400 spectrofluorometer (T-format) equipped with a rapid Peltier temperature controlled four-position sample holder with a GG495 longpass filter mounted. To start the reactions, 0.1 µM synaptobrevin (1-96) L26C-Alexa-488 was mixed with 1 µM SNAP-25 R136C-TMR and 1 µM syntaxin-1 (2-253)/Munc18-1 complexes (preformed for 1 hour at room temperature). The reaction buffer was 25 mM HEPES of pH 7.4, 150 mM KCl, 1 mM TCEP and 10% glycerol (v/v).

**Trans-SNARE complex formation assay.** Single cysteine variants of syntaxin-1A S186C and synaptobrevin-2 L26C were labeled with TMR and Alexa-488, respectively, as described above. V-liposomes were prepared similarly to those used for content mixing fusion assays (see below) except that they contained synaptobrevin-2 L26C-Alexa-488 with a 1:10,000 protein-to-lipid ratio. S-liposomes were prepared as the T-liposomes used for the content mixing assays but using syntaxin S186C-TMR without SNAP-25. Trans-SNARE complex formation was measured by the development of FRET between Alexa488-synaptobrevin on V- liposomes (0.0625 mM total lipid) and TMR-syntaxin-1A on S-liposomes (0.25 mM total lipid) at 37 °C using a PTI Quantamaster 400 spectrofluorometer (T-format) equipped with a rapid Peltier temperature controlled four-position sample holder. Prior to the measurements, the S-liposomes were incubated with 0.37 µM WT or mutant Munc18-1 for 1 hour at room temperature. The Alexa488 donor fluorescence of V-liposomes at 518 nm (excitation at 468 nm) was recorded to monitor the development of FRET over time upon mixing with S-liposomes containing the preformed syntaxin-1/Munc18-1 complexes, 2 µM SNAP-25a M71D, L78D, 0.2 µM Munc13C and 0.1 mM EGTA. At 1,100s, the reaction was paused and 0.6 mM $CaCl_2$ was quickly added to each reaction. The reaction buffer contained 25 mM HEPES of pH 7.4, 150 mM KCl, 1 mM TCEP and 10% glycerol (v/v). A GG495 long-pass filter (Edmund optics) was used to filter scattered light.

**Content mixing assays.** The assay was performed as previously described (*17*). Briefly, V-liposomes containing full-length synaptobrevin-2 (protein-to-lipid ratio 1:500) were prepared with 39% 1-palmitoyl, 2-oleoyl phosphatidylcholine (POPC), 19% 1,2-dioleoyl-sn-glycero-3-phospho-L-serine (DOPS), 19% 1-palmitoyl-2-oleoyl-sn-phosphatidylethanolamine (POPE), 20% cholesterol, 1.5% N-(7-nitrobenz-2-oxa-1,3-diazol-4-yl)-1,2-dihexadecanoyl-sn-glycero-3-phosphoethanolamine, triethylammonium salt (NBD-PE), and 1.5% 1,2-dihexadecanoyl-sn-glycero-3-phosphoethanolamine (Marina Blue DHPE). T-liposomes containing syntaxin-1 (WT or M183, or D184P, syntaxin-1-to-lipid ratio 1:800) and 25 µM SNAP-25 were prepared with 38% POPC, 18% DOPS, 20% POPE, 20% cholesterol, 2% phosphatidylinositol 4,5-biphosphate (PIP2), and 2% diacylglycerol (DAG). Dried lipid films were re-suspended in 25 mM HEPES, pH 7.4, 150 mM KCl, 1mM TCEP, 2% βOG. Lipid solutions were then mixed with the respective proteins and with 4 µM Phycoerythrin-Biotin for T-liposomes or with 8 µM Cy5-Streptavidin for V-liposomes in 25 mM HEPES, pH 7.4, 150mM KCl, 1mM TCEP, 10% glycerol (v/v). Proteoliposomes were prepared by detergent removal using dialysis with 2g/L Amberlite XAD-2 beads (Sigma) 3 times at 4 °C and subsequent co-floatation on a three-layer histodenz gradient (35%, 25%, and 0%), and harvested from the topmost layer. Content mixing signals were measured from the development of FRET between Cy5-streptavidin trapped in V-liposomes and Phycoerythrin-Biotin trapped in T-liposomes (excitation at 565 nm, emission at 670 nm). Each reaction was prepared in a total volume of 200 µl with V-liposomes (0.125 mM total lipid), T-liposomes (0.25 mM total lipids), 2.5 mM $MgCl_2$, 2 mM ATP, 0.1 mM EGTA, 5 µM streptavidin and the following proteins: 1 µM SNAP-25, 0.4 µM NSF purified in ATP containing buffer, 2 µM αSNAP, 1 µM Munc18-1 (WT or other tested mutants), 0.2 µM Munc13C. At 300 s, the reaction was paused and 0.6 mM $CaCl_2$ was added to each reaction mixture and quickly mixed. All experiments were performed at 30 ˚C using a PTI Quantamaster 400 spectrofluorometer (T-format) equipped with a rapid Peltier temperature controlled four-position sample holder. Content mixing measurements were normalized to control reactions collected without streptavidin in the presence of 1% βOG to measure the maximal Cy5 fluorescence attainable.

**Statistics.** To analyze the statistical significance of the results obtained in various assays (Figs. 3-5), all experiments were performed at least in triplicates and one-way analysis of variance (ANOVA) was performed through an all pairwise multiple comparison procedure using the Holm-Sidak test (*** $p < 0.001$, ** $p < 0.01$). For the analysis of binding of WT and Munc18-1 mutants to SyxLE/Syb by mass photometry (Fig. 3I), the difference between the $K_D$s obtained for WT and S42Q mutant Munc18-1 was obviously statistically significant ($p < 0.001$) but the very large $K_D$ calculated for the S42Q mutant and the correspondingly large standard deviation hinder the statistical analysis of significance for other pairwise comparisons within the group. Hence, all other comparisons within the group were performed after removing the S42Q data.

**Supplementary text**

It is now well established that the pathway that leads to neuronal SNARE complex assembly starts with closed syntaxin-1 tightly bound to Munc18-1 and proceeds through a template complex where both syntaxin-1 and synaptobrevin bind to Munc18-1. The binary Munc18-1-syntaxin-1 complex and the ternary synaptobrevin-Munc18-1-syntaxin-1 complexes contain extensive Munc18-1-SNARE interactions that are expected to confer specificity to SNARE complex formation but need to be released for the SNARE complex to form. Hence, the transition from the Munc18-1-syntaxin-1 complex to the SNARE complex must involve a series of steps where distinct inhibitory interactions are released, new interactions are formed, and some of the new and old interactions are released later. Interactions of the syntaxin-1 N-peptide and the $H_{abc}$ domain with Munc18-1 (Figs. 1D-F, S4A-C) are likely present throughout the pathway and may remain even after the SNARE complex is formed. It seems clear that these interactions, together with binding of the C-terminal half of the syntaxin-1 SNARE motif to the Munc18-1 cavity (Figs. 1D-F, S4A-C), are critical to keep syntaxin-1 bound to Munc18-1 while the N-terminal half of the SNARE motif and the linker undergo conformational rearrangements that lead to formation of the template complex. These rearrangements likely involve some changes in the orientation of the $H_{abc}$ domain with respect to Munc18-1 (Fig. S4D) but the $H_{abc}$ domain-Munc18-1 interface is similar in the Munc18-1-syntaxin-1 complex, class1 and class2 (Fig. S7D-F), suggesting that this interface is adaptable and can accommodate changes in relative orientation.

Unfurling the Munc18-1 loop to allow synaptobrevin binding to the groove formed by helices H11 and H12 (Fig. 2A-C) is undoubtedly a key event for SNARE complex assembly, as shown by the gain-of-function in vivo caused by the D326K mutation designed to unfurled the loop (*29*) and the enhancements of trans-SNARE complex assembly and of liposome fusion in the absence of $Ca^{2+}$ caused by this mutation (Figs. 4F and 5B). However, this mutation did not affect SNARE complex assembly in our solution assay (Fig. 4C). These observations suggests that synaptobrevin binding is a rate-limiting step in trans-SNARE complex assembly but not in the solution assays, which in principle is surprising because bridging of the two membranes by Munc13-1C (*14*) should increase the effective local concentration of synaptobrevin and the Munc18-1-syntaxin-1 complex between the membranes in the trans-SNARE complex assembly assay. Thus, it appears that in this assay Munc13-1C hinders binding of synaptobrevin to the Munc18-1-syntaixn-1 complex in the absence of $Ca^{2+}$, which likely arises because $Ca^{2+}$-free Munc13-1C has a tendency to bridge the membranes in an approximately perpendicular orientation that keeps the membranes apart (*14, 37*). It is also important to note that the P335A mutation in Munc18-1, which also induces a gain-of-function in vivo (*16, 25*), enhanced trans-SNARE complex assembly and liposome fusion (Figs. 4F, 5B), similar to the D326K mutation, but, also stimulated SNARE complex assembly in solution even in the absence of the Munc13-1 MUN domain (Fig. 4C,D), unlike the D326K mutation. Because P335 is well packed against syntaxin-1 in the Munc18-1-syntaxin-1 complex but not in class1 and class2 (Fig. S6I-K) and, correspondingly, the P335A mutation markedly impairs binding of Munc18-1 to syntaxin-1 but not to SyxLE/Syb (Figs. 3, S8), the gain-of-function caused by this mutation likely arises because it helps to release the N-terminal half of the syntaxin-1 SNARE motif and thus helps to form the template complex. Since the D326K mutation also helps to form the template complex, the release of interactions involving D326 and P335 are likely to be closely related events that might in principle occur in a concerted fashion, but the different effects of the D326K and P335A mutation in the solution SNARE complex assembly assay suggest that unfurling of the Munc18-1 loop occurs before release of the interactions between Munc18-1 P335 and the syntaxin-1 SNARE motif.

Conformational changes in syntaxin-1 are not required for unfurling of the Munc18-1 loop to allow binding of the C-terminal half of the synaptobrevin SNARE motif, but are necessary for further progress toward template complex formation. It is unclear whether the N-terminal half of the synaptobrevin SNARE motif might bind to the unfurled loop, as observed in the Vps33-Nyv1 complex (Fig. S5A), but it seems most likely that such binding is transient if it occurs, as it prevents interactions between the N-termini of the synaptobrevin and syntaxin-1 SNARE motifs for initiation of SNARE complex assembly. Such interactions require extension of the synaptobrevin helix toward the N-terminus, extension of helix 12 of Munc18-1 and movement of the helix formed by the syntaxin-1 the SNARE motif, which is long and bent in the Munc18-1-syntaxin-1 complex and becomes straight and shorter in class1 and class2 (Fig. 1D-F). This motion of the syntaxin-1 SNARE motif releases some of its interactions with the $H_{abc}$ domain that define the closed conformation (Figs. 2D-F, S7H-J) and with the Munc18-1 region containing P335 and a β-hairpin (Figs. 2A-C, S6I-K). Since the syntaxin-1 linker is packed against the SNARE motif in the closed conformation (Fig. S7G), the motion of the SNARE motif must also be accompanied by structural changes in the syntaxin-1 linker. A substantial energy barrier most likely needs to be overcome for all these rearrangements to occur, and interactions between a few residues of synaptobrevin and syntaxin-1 may not be sufficient to overcome this barrier. Even if such interactions occur transiently, the system might simply revert to the closed syntaxin-1-Munc18-1 complex structure unless the initial synaptobrevin and syntaxin-1 interactions are stabilized.

The observation of the short four-helix bundle between helices Hd and He from the syntaxin-1 linker and the SNARE motifs of syntaxin-1 and synaptobrevin in both class1 and class2 (Figs. 1E,F, 2E,F), together with the severe impairment in liposome fusion caused by the M183E and D184P mutations in the linker (Fig. 5C,D), indicate that formation of this short four-helix bundle is a key step to trigger the dissociation of interactions involving the syntaxin-1 SNARE motif necessary to initiate SNARE complex assembly. It is noteworthy that the linker forms a similar small four helix bundle with the syntaxin-1 and synaptobrevin SNARE motifs in both class1 and class2 (Fig 2J) despite considerable differences in the structures of class1 and class2 around this region, including the bend of the synaptobrevin helix in class2 (Figs. 1F, 2C) and the different orientations of the short four-helix bundle with respect to the $H_{abc}$ domain and the C-terminal half of the syntaxin-1 SNARE motif (Fig. 2E,F). These observations suggest that this small four-helix bundle constitutes a relatively stable (or metastable) structure that helps to nucleate the first interactions between syntaxin-1 and synaptobrevin for eventual SNARE complex formation. The less extensive contacts between the SNARE motif of syntaxin-1 and the $H_{abc}$ domain in class2 compared to class1 (Figs. 2E,F S7I,J) strongly suggest that class1 (and/or similar structures) occurs earlier than class2 (and/or similar structures) (Fig. 6C,D) and that the short-four helix bundle holds the SNARE motifs of syntaxin-1 and synaptobrevin SNARE motifs together while additional interactions that hinder progress to SNARE complex assembly are released. Note however that the structure of this short four-helix bundle is likely dynamic, as the electron density in this region was poor (Fig. S2B), and is necessarily transient as the linker needs to dissociate from the syntaxin-1 and synaptobrevin SNARE motifs to allow SNAP-25 binding and SNARE complex assembly.

Comparisons of class1 and class2 with the crystal structure of the yeast SM protein Vps45 bound to the syntaxin-1 homologue Tlg2 (*51*), which provides a reference for a syntaxin homologue in an open conformation, show that the orientation of the $H_{abc}$ domain of Tlg2 resembles more that observed in class1 than that of class2 (Fig. S5G-I). It is possible that the preferred orientation of the $H_{abc}$ domain from open syntaxin-1 with respect to Munc18-1 is different from that observed in the Vps45/Tlg2 complex and is similar to that observed in the closed syntaxin-Munc18-1 complex and in class2, but this orientation changes transiently when class1 is formed to allow extension of helix 12 of Munc18-1 and to facilitate interactions between the N-termini of the synaptobrevin and syntaxin-1 SNARE motifs; subsequently, the $H_{abc}$ domain/Munc18-1 interface may relax back to the preferred orientation as syntaxin-1 opens more to form class2, with concomitant bending of the synaptobrevin helix (Fig. 6B-D). It is also plausible that the different elements that need to be re-arranged to form the template complex and eventually the SNARE complex may go back and forth between distinct positions in a less coordinated fashion until a sufficient number of productive events coincide to overcome the energy barriers hindering template complex formation.

It is also important to note that Munc13-1 is expected to influence the molecular events outlined above by at least two mechanisms. First, as mentioned above, the orientation of Munc13-1 as it bridges the two membranes may control the accessibility of the Munc18-1-syntaxin-1 complex to synaptobrevin. Second, the Munc13-1 MUN domain helps open syntaxin-1 through weak interactions with a region of the syntaxin-1 linker (residues 150-160) (*39, 40*) that is likely flexible, as it was not observed in our cryo-EM structures. These interactions might facilitate formation of the small four-helix bundle involving the linker and the syntaxin-1 and synaptobrevin SNARE motifs, or may simply destabilize the linker conformation existing in the Munc18-1-syntaxin-1 complex, thus catalyzing conformational rearrangements in syntaxin-1. Both effects or perhaps direct Munc18-1-Munc13-1 interactions might also stimulate unfurling of the Munc18-1 loop to facilitate synaptobrevin binding. In this scenario, Munc13-1 binding to the linker would be the first event, followed by loop unfurling and synaptobrevin binding. It is also plausible that loop unfurling and synaptobrevin binding occurs first, as proposed above, but progress toward template complex formation can occur only when Munc13-1 binds to the linker, facilitating conformational rearrangements in syntaxin-1. Munc13-1 was also proposed to facilitate SNARE complex assembly through interactions with helix 11 of Munc18-1 involving Q301 (*34*), although our mass photometry data (Fig. 3J) support the alternative hypothesis that the functional effects of mutating Q301 arise because of disruption of Munc18-1-synaptobrevin binding (*35*).

While some of the details of this model are still unclear, the overall model readily explains the gains-of-function caused by the D326K and P335A mutations, as outlined above, and also the impairments in trans-SNARE complex assembly and liposome fusion caused by the Q301D, L307R, L348R and E352K mutations, which can be attributed to the disruption of synaptobrevin binding caused by these mutations (Figs. 4F, 5B). The observation that the Q301D, L307R, L348R and E352K mutations did not impair SNARE complex assembly in solution (Fig. 4C) reinforces the conclusion that synaptobrevin binding to Munc18-1 is not rate limiting in these assays.

The effects of the S42Q mutation in our assays were less consistent. The facilitation of SNARE complex assembly in solution and of liposome fusion caused by this mutation (Figs. 4C,D and 5B) suggest that release of the interactions between the C-terminal half of the syntaxin-1 SNARE motif and Munc18-1 (Figs. 1D-F, S7A-C) to allow final SNARE complex zippering and fusion constitutes one of the rate-limiting steps in these assays. However, the S42Q mutation impaired trans-SNARE complex assembly (Fig. 4F). The basis for these differential effects is unclear, but it is most likely that they arise because of differences in the conditions of the experiments and/or in the corresponding rate limiting steps. For instance, it is plausible that the S42Q mutation indeed stimulates liposome fusion because it facilitates release of the interaction of Munc18-1 with the C-terminal half of the syntaxin-1 SNARE motif, whereas Munc18-1 may remain bound to part of the C-terminal half of the syntaxin-1 SNARE motif in the partially formed SNARE complex resulting in the trans-SNARE complex assembly assays [note that these assays were performed with SNAP-25 bearing mutations that hinder final C-terminal zippering of the four-helix bundle to prevent fusion (*10*)]. In this case, the slower rate of trans-SNARE complex assembly observed with the S42Q Munc18-1 mutant may arise because the mutation destabilizes the transition state in this reaction more severely than the initial syntaxin-1-Munc18-1 complex. It is also possible that the rate limiting steps of the trans-SNARE complex assembly assays and liposome fusion assays are different because of the different synaptobrevin-to-lipid ratios used in these assays (1:10,000 and 1:500, respectively) and that the different effects of the S42Q mutation in the two assays arise because the mutation causes differential destabilization of the corresponding transition states. Regardless of these possibilities, the differences in the effects of the S42Q and P335A mutations in trans-SNARE complex assembly indicate that the interactions with syntaxin-1 involving the mutated residues are released at different stages, as expected from the general belief that the SNARE complex assembles from the N-terminus to the C-terminus. Overall, the effects of the different mutations that we used in this study vividly illustrate how multiple energy barriers hinder the transitions that lead from the Munc18-1-syntaxin-1 complex to the SNARE complex.

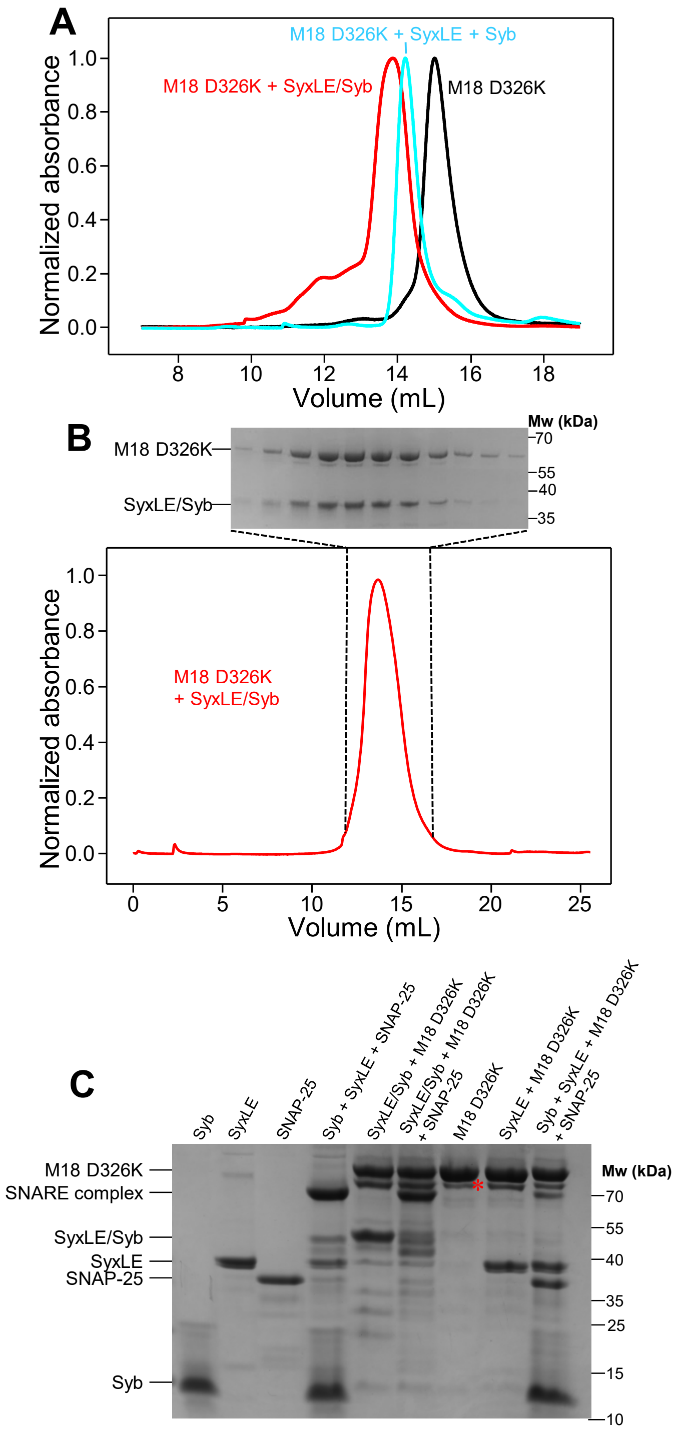


**Fig. S1.** Preparation of the template complex. (**A**) Gel filtration on Superdex S200 of Munc18-1 D326K alone (black curve) or after overnight incubation with SyxLE/Syb (red curve) or after overnight incubation with non-cross-linked SyxLE plus Syb (blue curve). The absorbance at 280 nm was normalized to the maximum absorbance observed in each chromatogram. (**B**) Gel filtration of an equimolar mixture of Munc18-1 D326K and SyxLE/Syb used to purify the template complex. SDS-PAGE analysis of the eluted fractions containing the complex is shown above the chromatogram. (**C**) SDS-PAGE analysis of mixtures of Syb, SyxLE, SNAP-25, Munc18-1 D326K (M18) and SyxLE/Syb in different combinations as indicated. All samples were incubated for three minutes at room temperature before loading onto the gel. Formation of the SDS-resistant SNARE complex was much more efficient for the mixture containing SyxLE/Syb. The positions of the proteins and complexes are indicated on the left, and those of molecular weight markers on the right. A degradation band of Munc18-1 is indicated with a *, but note that such degradation is common in Munc18-1 and does not prevent syntaxin-1 binding (*8*).

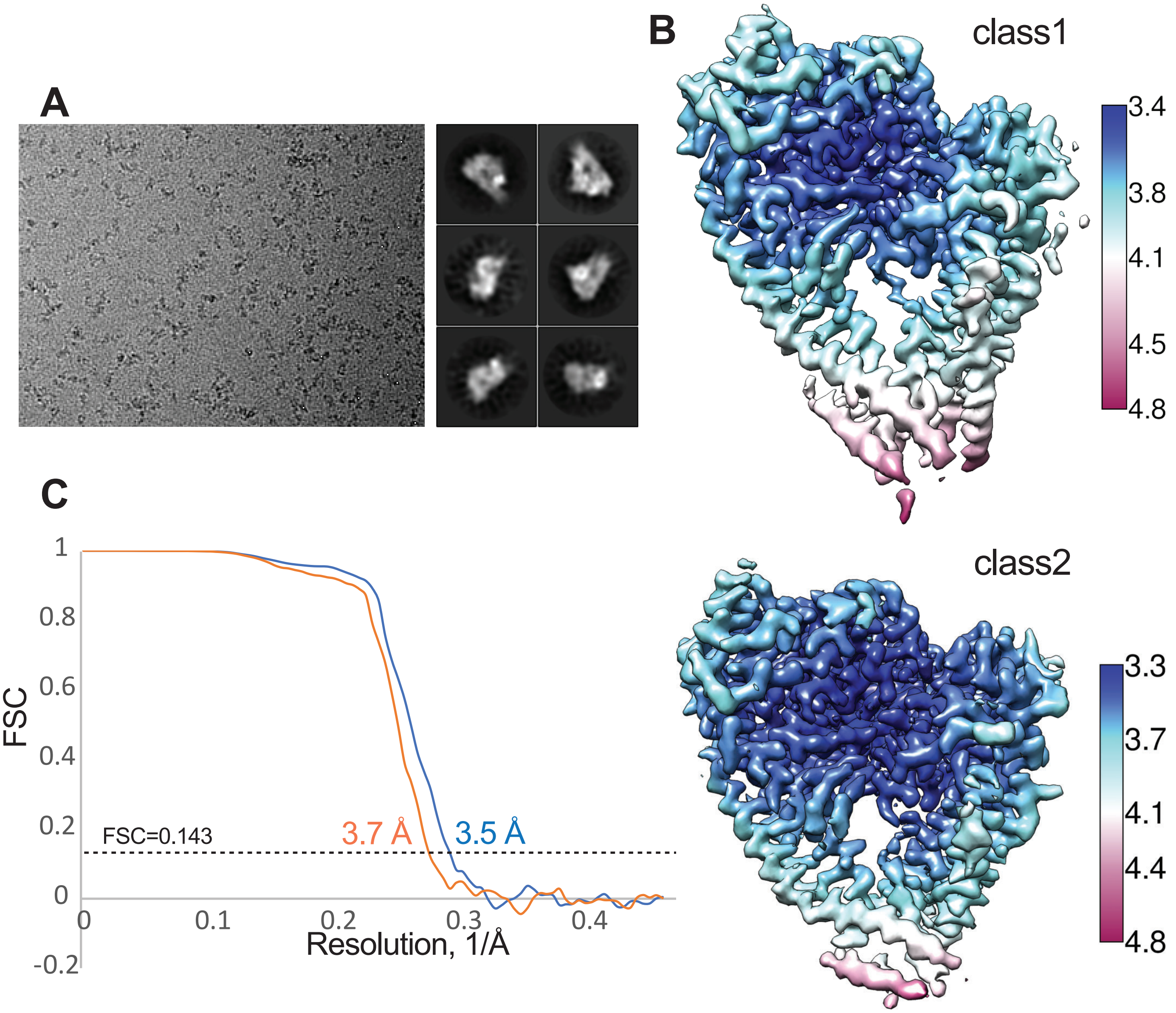


**Fig. S2.** Cryo-EM analysis of the template complex. (**A**) Representative electron micrograph from a total 7,401 images acquired and 2D class averages of the template complex. (**B**) Cryo-EM maps of the two conformers of the template complex (class1 and class2) colored by local resolution. (**C**) Gold-standard Fourier shell correlation (FSC) curve for the cryo-EM maps of the template complex. Red: class1; blue: class2.

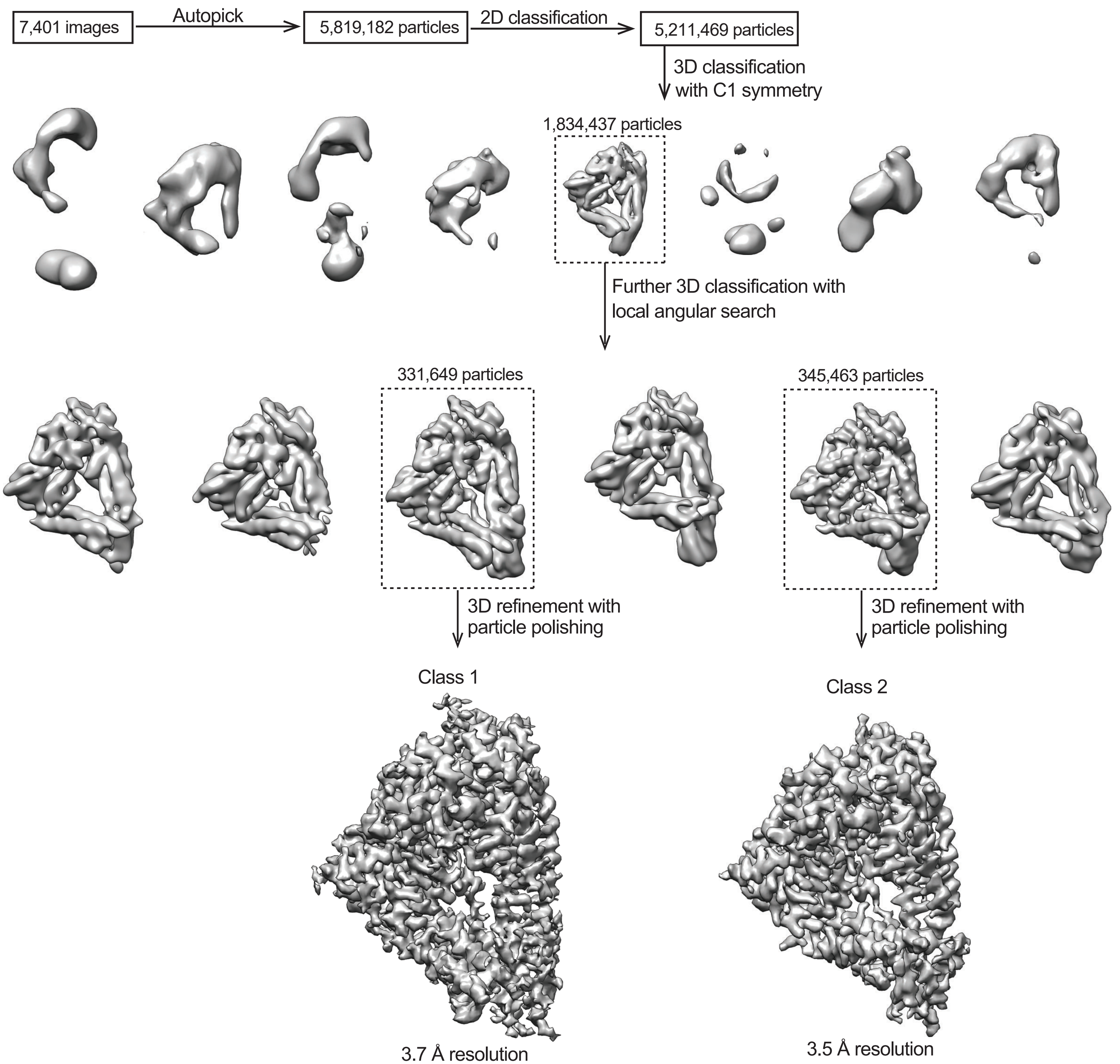


**Figure S3.** Flow-chart of cryo-EM data processing leading to the two conformers of the template complex (class1 and class2).

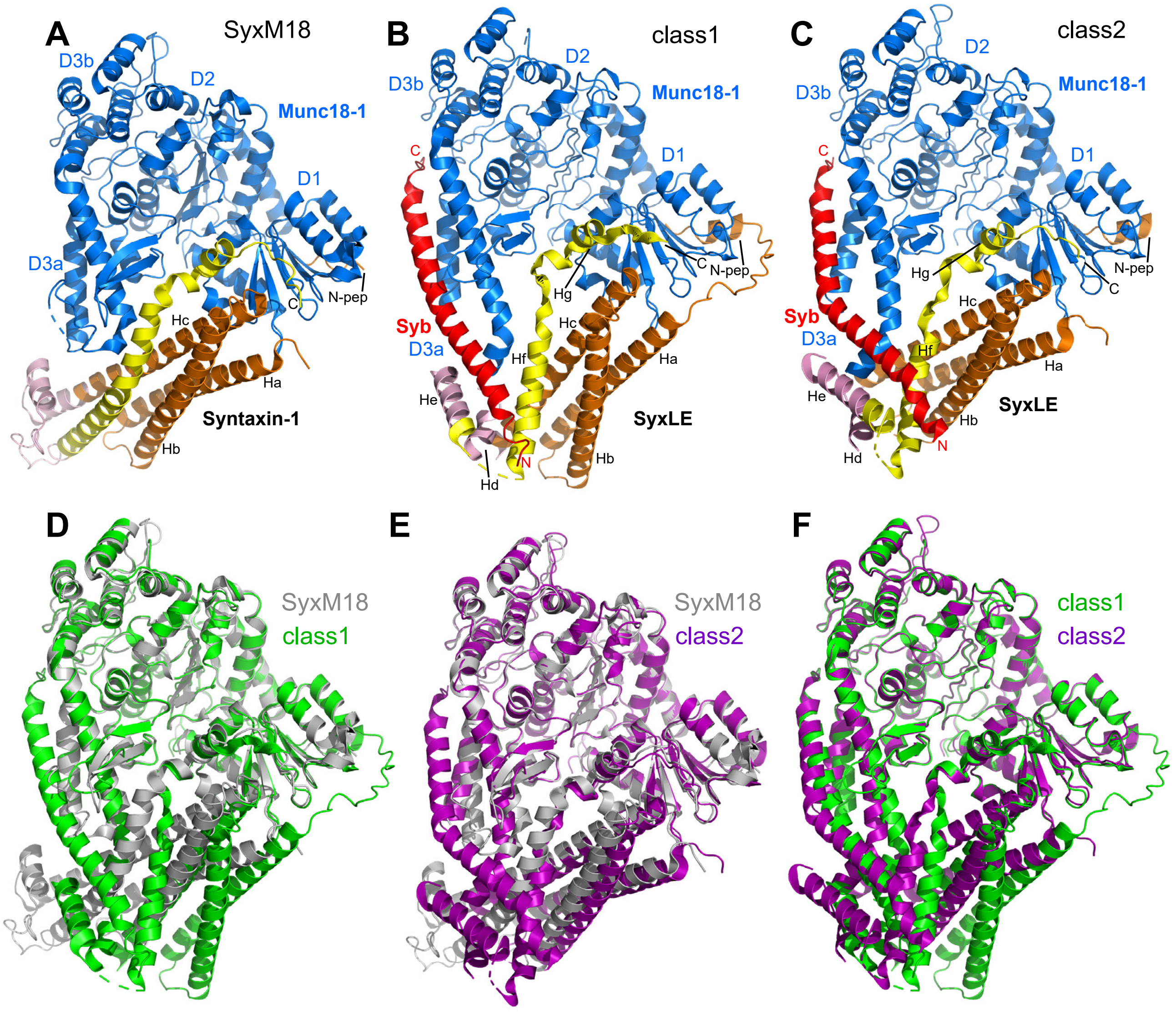


**Figure S4.** Structural comparison of class1, class2 and the syntaxin-1-Munc18-1 complex (SyxM18). (**A-C**) Ribbon diagrams of the structures of the syntaxin-1-Munc18-1 complex (SyxM18) (PDB code 3C98) (*8, 32*) (**A**), class1 (**B**) and class2 (**C**). The color code is the same as in Fig. 1D-F. The domains of Munc18-1 (D1, D2, D3a and D3b) are labeled. The helices formed by syntaxin-1 in class1 and class2 (named Ha-Hg) are indicated. Helices Ha-Hc are also present in the syntaxin-1-Munc18-1 complex, but the other helices are remodeled. (**D-F**) Superpositions of ribbon diagrams of SyxM18 (grey), class1 (green) and class2 (purple) using the Munc18-1 C$\alpha$ atoms for the superpositions. The root mean square (r.m.s.) deviations were the following: Munc18-1 versus class1 (**D**) 0.59 Å for 470 C$\alpha$ carbons; Munc18-1 versus class2 (**E**) 0.54 Å for 455 C$\alpha$ carbons; and class1 versus class2 (**F**) 0.38 Å for 474 C$\alpha$ carbons.

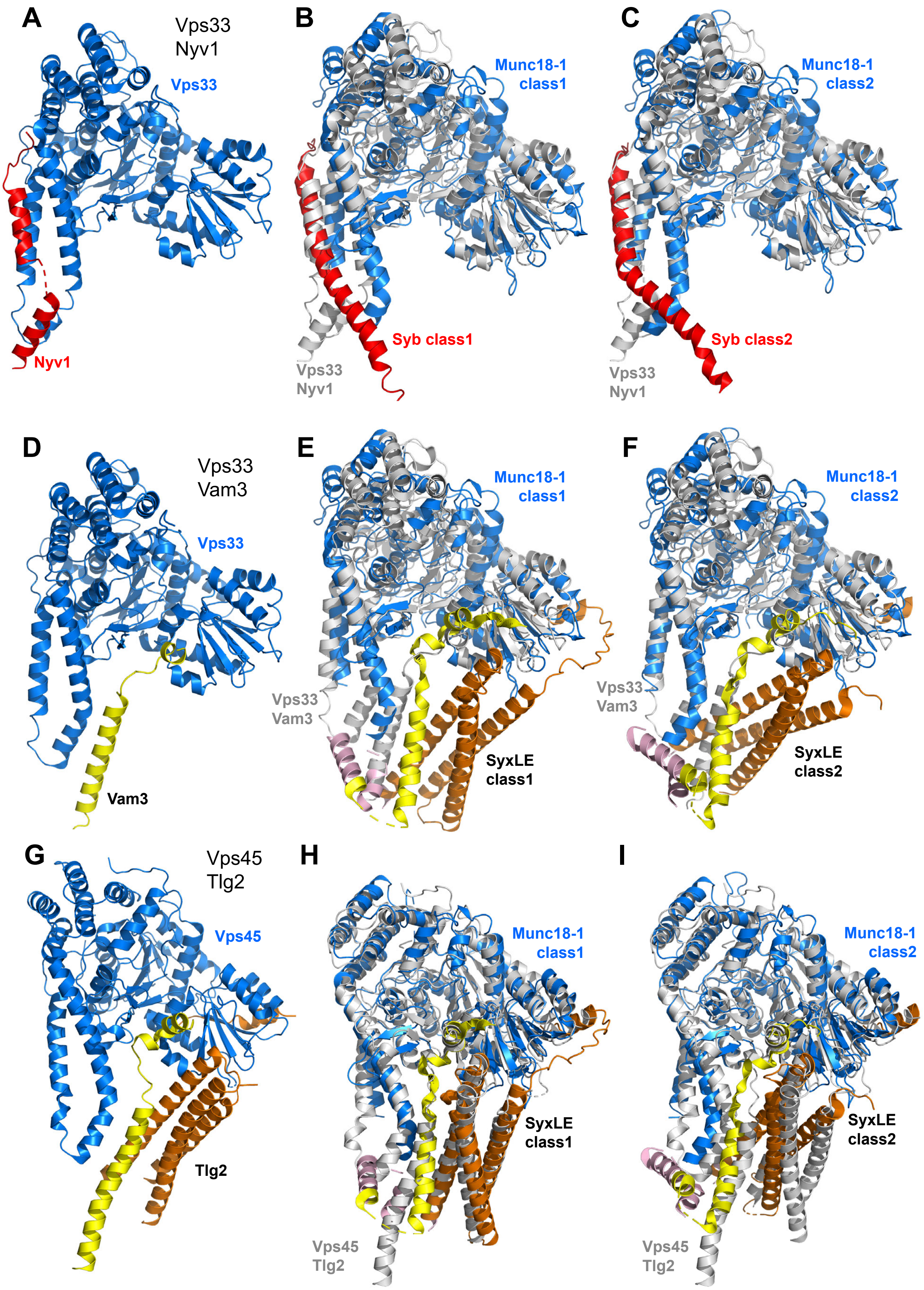

A
Vps33
Nyv1
Vps33
Nyv1
B
Munc18-1
class1
Syb class1
Vps33
Nyv1
C
Munc18-1
class2
Syb class2
Vps33
Nyv1
D
Vps33
Vam3
Vps33
Vam3
E
Munc18-1
class1
Vps33
Vam3
SyxLE
class1
F
Munc18-1
class2
Vps33
Vam3
SyxLE
class2
G
Vps45
Tlg2
Vps45
Tlg2
H
Munc18-1
class1
SyxLE
class1
Vps45
Tlg2
I
Munc18-1
class2
SyxLE
class2
Vps45
Tlg2

**Figure S5.** Structural comparisons of class1 and class 2 with the Vps33/Nyv1, Vpss33/Vam3 and Vps45/Tlg2 complexes. (**A**,**D**,**G)** Ribbon diagrams of the structures of the Vps33-Nyv1 complex (PDB code 5BV0) (*28*) (**A**), the Vps33-Vam3 complex (PDB code 5BUZ) (*28*) (**D**) and the Vps45-Tlg complex (PDB code 6XMD) (*51*) (**G**). Note that the Vps33-Nyv1 and Vps33-Vam3 complexes also included Vps16, which is not shown for simplicity. Vps33 and Vps45 are shown in blue, Nyv1 in red, the Vam3 SNARE motif in yellow, and Tlg2 in orange ($H_{abc}$ domain) and yellow (SNARE motif). (**B**,**C**,**E**,**F**,**H**,**I**) Superpositions of class1 and class2 with Vps33-Nyv1 (**B**,**C**), Vps33-Vam3 (**E**,**F**) and Vps45-Tlg2 (**H**,**I**). Vps33-Nyv1, Vps33-Vam3 and Vps45-Tlg2 are shown in gray. The color code for class1 and class 2 is the same as in Fig. 1. The superpositions per performed using corresponding C$\alpha$ carbons of the SM proteins, yielding the following r.m.s. deviations: Vps33-Nyv1 versus class1: 4.91 Å for 394 C$\alpha$ carbons; Vps33-Nyv1 versus class2: 5.18 Å for 411 C$\alpha$ carbons; Vps33-Vam3 versus class1 4.47 Å for 379 C$\alpha$ carbons; Vps33-Vam3 versus class2 4.99 Å for 406 C$\alpha$ carbons; Vps45-Tlg2 versus class1 1.67 Å for 323 C$\alpha$ carbons; and Vps45-Tlg2 versus class2 1.80 Å for 323 C$\alpha$ carbons.

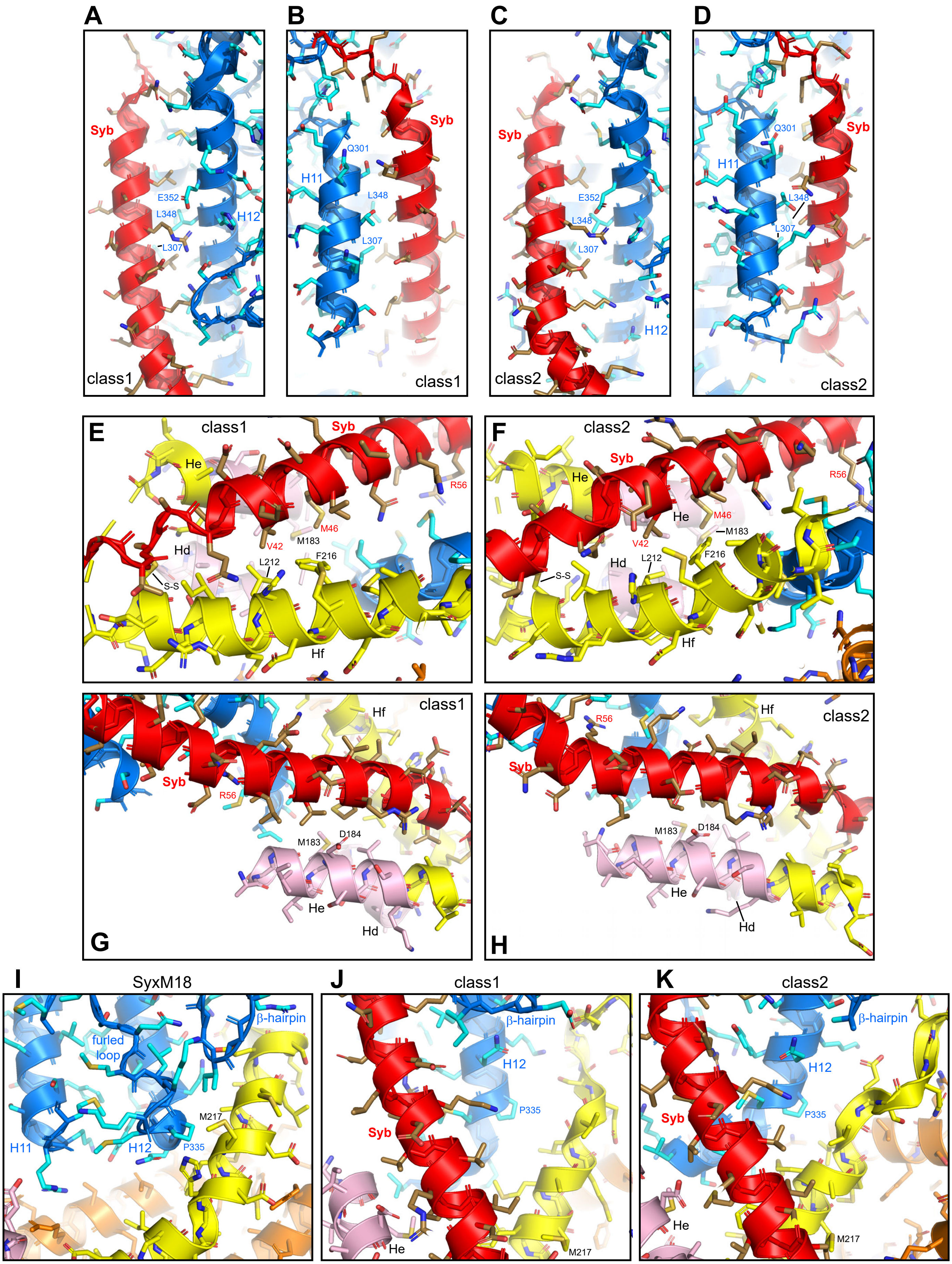

A
Syb
E352
L348
L307
H12
class1
B
Syb
Q301
H11
L348
L307
class1
C
Syb
E352
L348
L307
H12
class2
D
Syb
Q301
H11
L348
L307
class2
E
class1
Syb
He
R56
M46
V42
M183
Hd
L212
F216
S-S
Hf
F
class2
Syb
He
R56
M46
M183
V42
Hd
L212
F216
S-S
Hf
G
class1
Hf
Syb
R56
M183
D184
He
Hd
H
class2
R56
Hf
Syb
M183
D184
He
Hd
I
SyxM18
β-hairpin
furled loop
M217
H11
H12
P335
J
class1
β-hairpin
H12
P335
Syb
He
M217
K
class2
β-hairpin
H12
Syb
P335
He
M217

**Figure S6.** Interactions between different structural elements of Munc18-1, synaptobrevin and syntaxin-1 in class1 and class2. (**A**-**K**) The diagrams show close-up views of the regions where: synaptobrevin binds to Munc18-1 in class1 (**A**,**B**) and class2 (**C**,**D**); the syntaxin-1 linker forms a small four-helix bundle with the syntaxin-1 and synaptobrevin SNARE motifs in class1 (**E**,**G**) or class2 (**F**,**H**); P335 and a β-hairpin of Munc18-1 interact with or are close to the syntaxin-1 motif in the syntaxin-1-Munc18-1 complex (SyxM18) (**I**), class1 (**J**) or class2 (**K**) The structures are represented with ribbon diagrams [Munc18-1 in blue; synaptobrevin (Syb) in red; syntaxin-1 in orange (N-peptide and $H_{abc}$ domain), pink (linker) and yellow (SNARE motif)] and stick models with the following color code for side chain atoms: oxygen atoms red, nitrogen atoms blue, sulfur atoms light orange and carbon atoms in cyan for Munc18-1, brown for synaptobrevin and orange ($H_{abc}$ domain), pink (linker) or yellow (SNARE motif) for syntaxin-1. The positions of selected residues discussed in the text, selected helices of syntaxin-1, the disulfide bond linking syntaxin-1 and synaptobrevin, the furled loop and the β-hairpin are indicated in the relevant panels. The M217 side chain is labeled in panels **I**-**K** to show that the SNARE motif of syntaxin-1 is shifted downwards in class1 and class2 with respect to its position in the syntaxin-1-Munc18-1 complex.

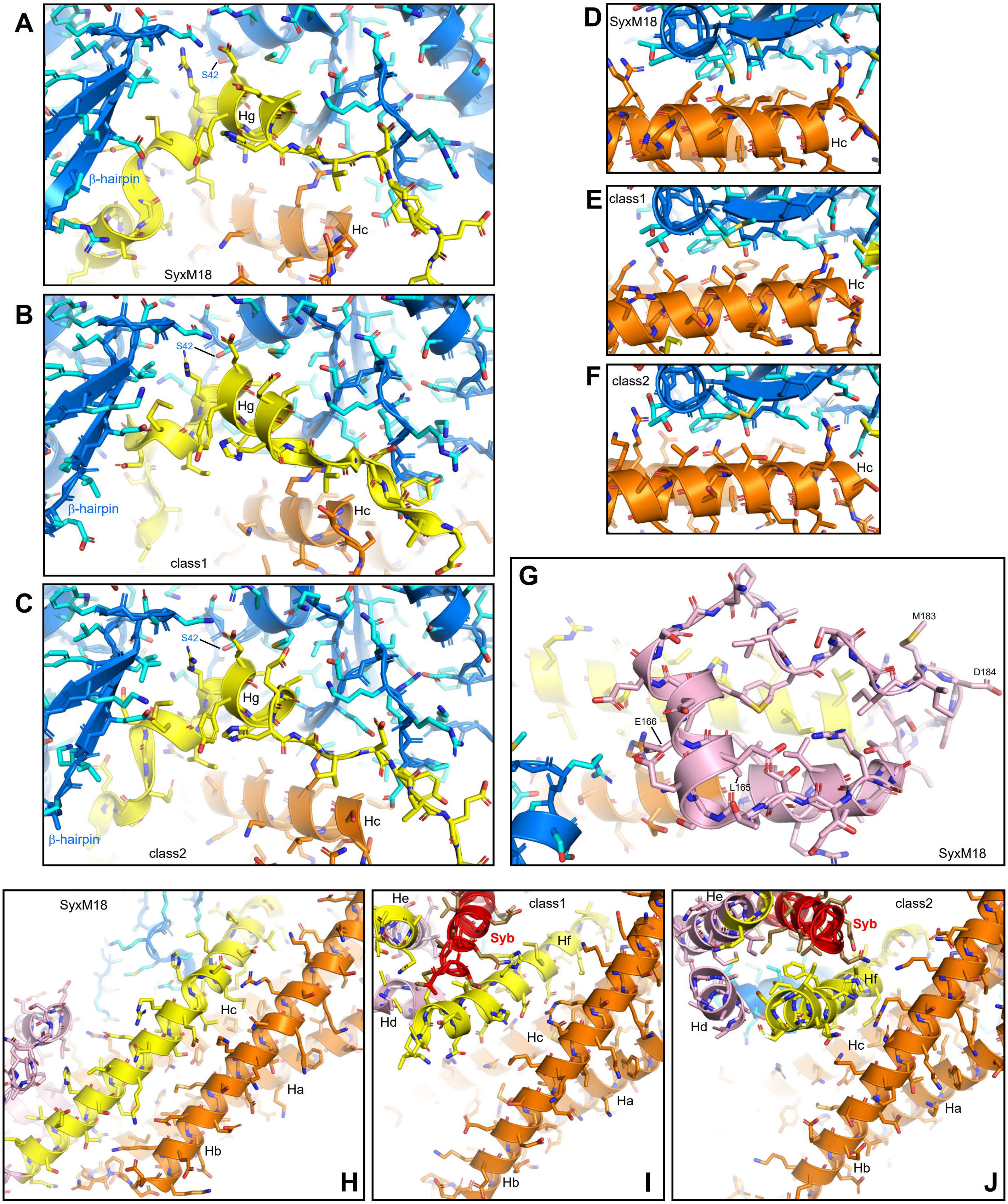

A
S42
Hg
β-hairpin
Hc
SyxM18
B
S42
Hg
β-hairpin
Hc
class1
C
S42
Hg
β-hairpin
class2
Hc
D
SyxM18
Hc
E
class1
Hc
F
class2
Hc
G
M183
D184
E166
L165
SyxM18
SyxM18
Hc
Ha
Hb
H
He
class1
Syb
Hf
Hd
Hc
Ha
Hb
I
He
class2
Syb
Hf
Hd
Hc
Ha
Hb
J

**Figure S7.** Comparison of interactions among Munc18-1, the syntaxin-1 SNARE motif and the $H_{abc}$ domain in the syntaxin-1-Munc18-1 (SyxM18) complex, class1 and class2. The diagrams show cloase-up views of the regions where: the C-terminal half of the syntaxin-1 SNARE motif binds to the Munc18-1 cavity in the syntaxin-1-Munc18-1 complex (SyxM18) (**A**), class1 (**B**) or class2 (**C**); the syntaxin-1 $H_{abc}$ domain binds to Munc18-1 in SyxM18 (**D**), class1 (**E**) or class2 (**F**); the syntaxin-1 linker packs against the SNARE motif in SyxM18 (**G**); and the syntaxin-1 SNARE motif interacts with the $H_{abc}$ domain in SyxM18 (**H**), class1 (**I**) or class2 (**J**). The structures are represented with ribbon diagrams and stick models with the same color coding as Fig. S6. The positions of selected helices and residues discussed in the text are indicated in the relevant panels. Panel (**G**) shows that residues L165 and E166, which are replaced in the LE mutation, are buried in SyxM18.

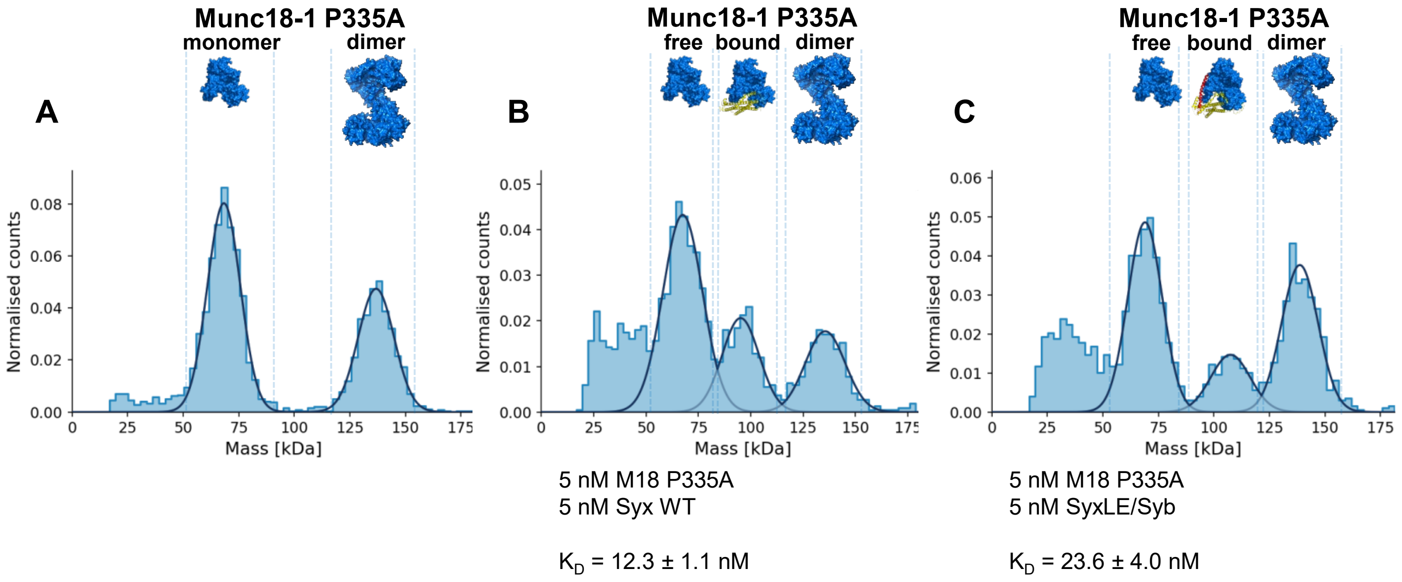


**Figure S8.** Analysis of Munc18-1 P335A mutant-SNARE interactions by mass photometry. (**A**-**C**) Normalized histograms of mass distributions observed for samples containing 5 nM Munc18-1 (M18) P335A mutant alone (**A**) or together with 5 nM WT syntaxin-1(2-253) (Syx) (**B**), or 5 nM SyxLE/Syb (**C**). Gaussian fits (solid lines) were used to calculate the populations of free and bound Munc18-1 P335A, and derive dissociation constants ($K_D$s).

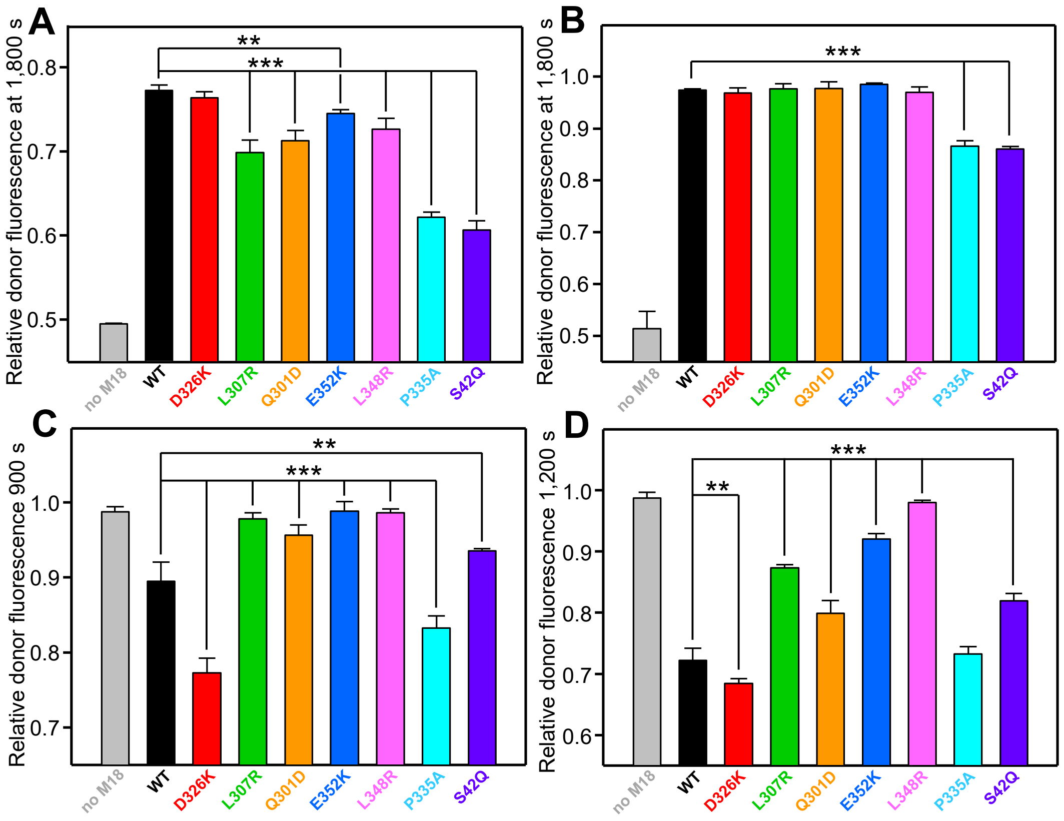


**Figure S9.** Quantification of the SNARE complex assembly assays at selected time points to highlight differences between the results obtained with different mutants. (**A**,**B**) Diagrams showing the relative donor fluorescence (normalized to the first time point) observed at 1,800 s in the SNARE complex assembly assays performed in solution with SyxMUN (**A**) or Syx (**B**) in the presence of WT or mutant Munc18-1 (corresponding to Fig. 4C and 4D, respectively). (**C**,**D**) Diagrams showing the relative donor fluorescence (normalized to the first time point) observed at 900 s, i.e. before $Ca^{2+}$ addition (**C**), and at 1,200 s, i.e. after $Ca^{2+}$ addition (**D**), in the trans-SNARE complex performed with WT or mutant Munc18-1 (corresponding to Fig. 4F). Bars represent averages of the normalized fluorescence intensities observed in three independent experiments. Error bars represent standard deviations. Statistical significance and p values were determined by one-way analysis of variance (ANOVA) with the Holm-Sidak test (*** $p < 0.001$, ** $p < 0.01$).

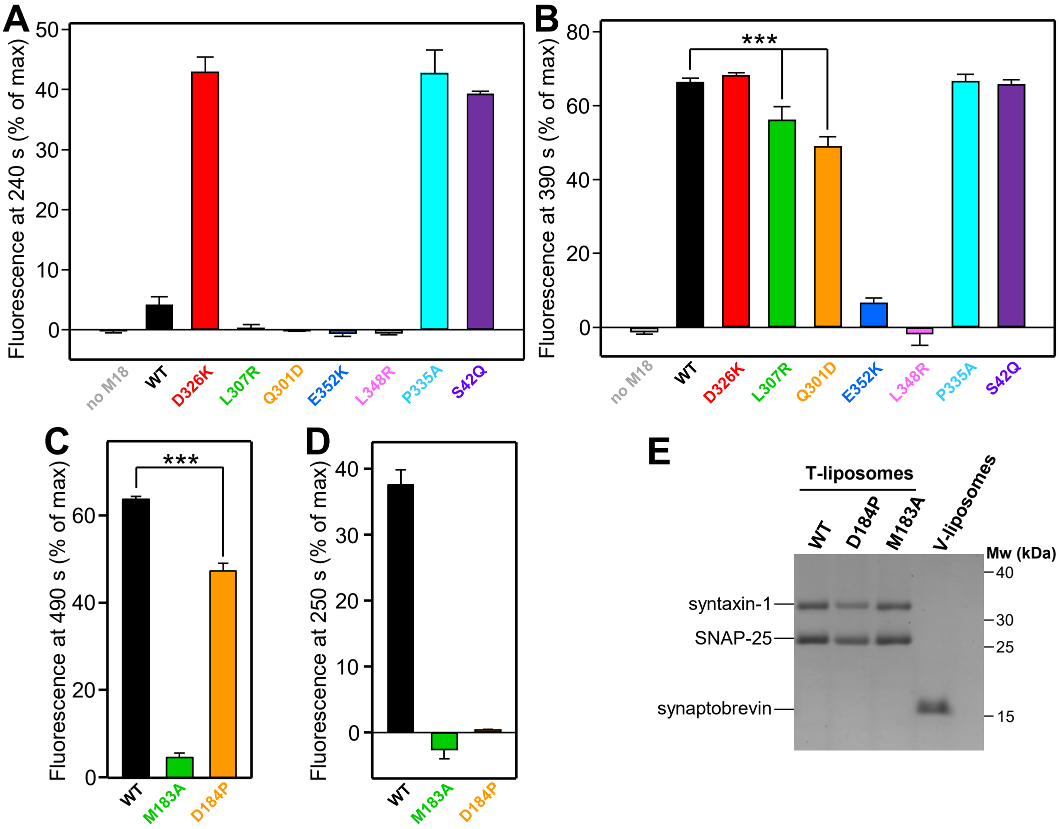


**Figure S10.** Quantification of the liposome fusion assays performed with different Munc18-1 and syntaxin-1 mutants. (**A**-**D**) Diagrams showing the relative fluorescence observed at 240 s, i.e. before $Ca^{2+}$ addition (**A**), and at 390 s, i.e. after $Ca^{2+}$ addition (**B**), in the content mixing assays performed with WT and mutant Munc18 (corresponding to Fig. 5B); at 490 s, i.e. after $Ca^{2+}$ addition in the content mixing assays performed with WT Munc18-1 and WT or mutant syntaxin-1 (**C**) (corresponding to Fig. 5C); or at 250 s, i.e. before $Ca^{2+}$ addition, in analogous assays performed with D326K mutant Munc18-1 and WT or mutant syntaxin-1 (**D**) (corresponding to Fig. 5D). Bars represent averages of the normalized fluorescence intensities observed in three independent experiments. Error bars represent standard deviations. Statistical significance and p values were determined by one-way analysis of variance (ANOVA) with the Holm-Sidak test (*** $p < 0.001$). (**E**) SDS-PAGE analysis of protein incorporation into V-liposomes and the T-liposomes containing WT or mutant syntaxin-1, which were used for the fusion assays of Fig. 5. The positions of the SNARE proteins are indicated on the left, and those of molecular weight markers on the right.

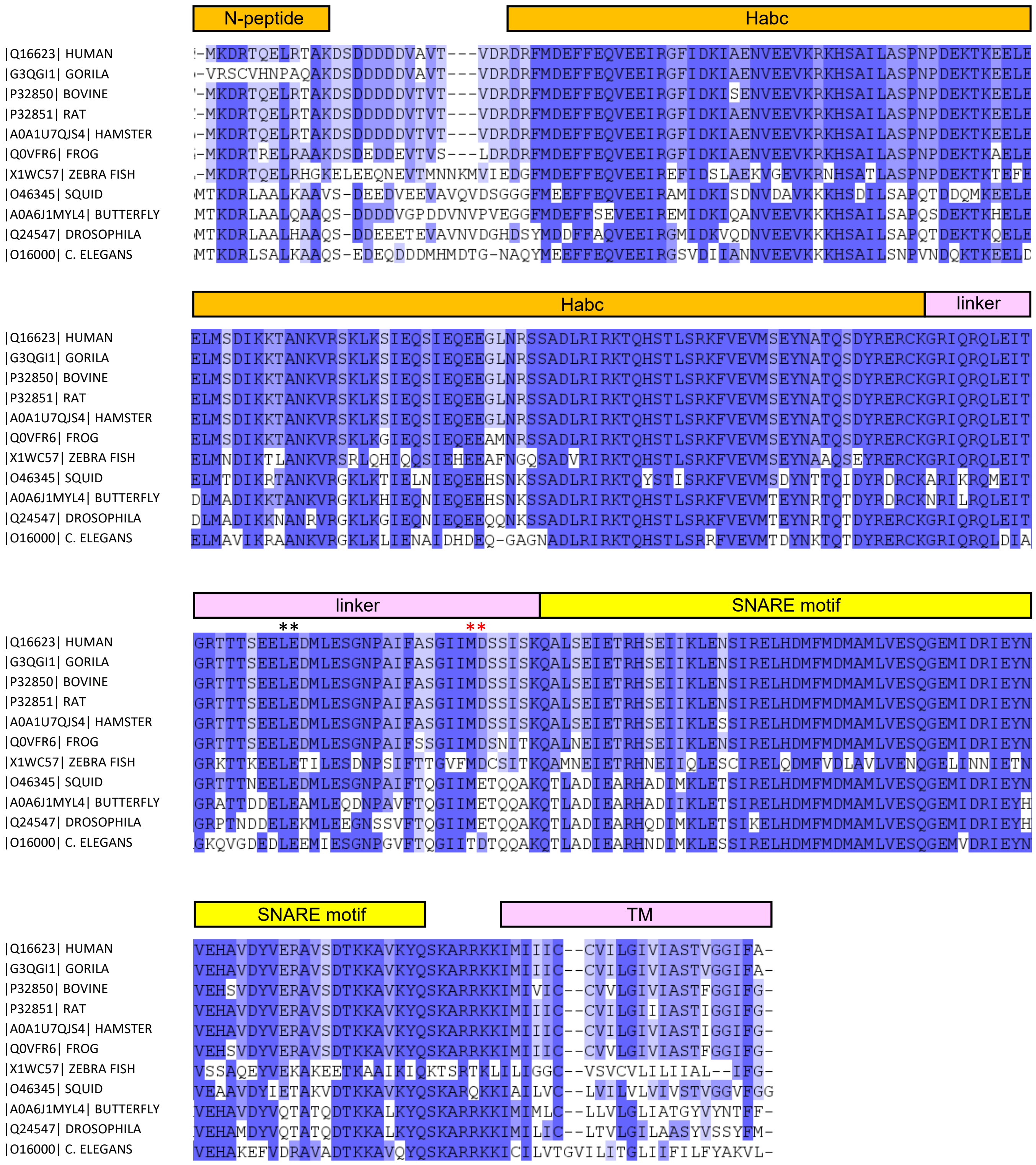


**Figure S11.** Sequence alignment of syntaxin-1A homologues from different species. The Uniprot (https://www.uniprot.org) code of each sequence and the common name of the species are indicated on the left. Conserved residues have a colored background. The color intensity reflects the degree of conservation. The domain diagram of syntaxin-1 with the same color code used in Fig. 1A is shown above the sequences. The positions of L165 and L166, which were mutated in the syntaxin-1 LE mutant, and of M183 and D184, the residues in the linker that were mutated, are indicated with *.

**Supplementary Tables**

**Table S1. Cryo-EM data collection, refinement, and validation statistics.**

| | class2 | class1 |
|---|---|---|
| **Data collection and processing** | | |
| Magnification | 46,296 | 46,296 |
| Voltage (kV) | 300 | 300 |
| Electron exposure (e–/Å2) | 60 | 60 |
| Defocus range (μm) | 1.6 – 2.6 | 1.6 – 2.6 |
| Pixel size (Å) | 1.08 | 1.08 |
| Symmetry imposed | C1 | C1 |
| Initial particle images (no.) | 5,819,182 | 5,819,182 |
| Final particle images (no.) | 345,463 | 331,649 |
| Map resolution (Å) | 3.5 | 3.7 |
| FSC threshold | 0.143 | 0.143 |
| **Refinement** | | |
| Initial model used (PDB code) | 3C98 | 3C98 |
| Model composition | | |
| Atoms | 6464 | 6425 |
| Protein residues | 806 | 806 |
| Ligands | 0 | 0 |
| R.m.s. deviations | | |
| Bond lengths (Å) | 0.003 | 0.003 |
| Bond angles (°) | 0.529 | 0.539 |
| **Validation** | | |
| MolProbity score | 1.53 | 1.63 |
| Clashscore | 6.39 | 7.77 |
| Poor rotamers (%) | 0.00 | 0.28 |
| Ramachandran plot | | |
| Favored (%) | 96.94 | 96.70 |
| Allowed (%) | 3.06 | 3.30 |
| Disallowed (%) | 0.00 | 0.00 |

**Table S2. Buried accessible surface areas (Å$^2$) between different proteins and structural elements in class1, class2 and the Munc18-1-syntaxin-1 complex**[a]

| | Munc18-1 syntaxin-1 | Munc18-1 synaptobrevin | Munc18-1 SyxLE/Syb | syntaxin-1 synaptobrevin | syntaxin-1 $H_{abc}$ domain syntaxin-1 SNARE motif |
|---|---|---|---|---|---|
| class1 | 5619 | 2825 | 8389 | 1320 | 1019 |
| class2 | 5487 | 2826 | 8267 | 1361 | 876 |
| SyxM18[b] | 5406 | | | | 2626 |

[a]Buried solvent accessible surface areas were calculated using Pymol (Schrödinger, Inc.) adding hydrogen atoms and using high sampling density, with the parameter dot_solvent set to 1 and set dot_density set to 3.

[b]Syntaxin-1-Munc18-1 complex (PDB code 3C98)

**Table S3. Summary of concentrations used for mass photometry experiments and measured $K_D$s[a]**

| Protein 1 | [Protein 1] | Protein 2 | [Protein 2] | $K_D$ (nM) |
|---|---|---|---|---|
| Munc18-1 WT | 5 nM, 5 nM | syntaxin-1 WT | 2.5 nM, 5 nM | 3.3 ± 0.5 |
| Munc18-1 P335A | 5 nM, 5 nM | syntaxin-1 WT | 2.5 nM, 5 nM | 12.3 ± 1.1 |
| Munc18-1 S42Q | 5 nM, 10 nM | syntaxin-1 WT | 5 nM, 5 nM | 67.9 ± 4.3 |
| Munc18-1 WT | 5 nM, 5 nM | syntaxin-1 M183A | 2.5 nM, 5 nM | 0.6 ± 0.2 |
| Munc18-1 WT | 5 nM, 5 nM | syntaxin-1 D184P | 2.5 nM, 5 nM | 1.9 ± 0.2 |
| Munc18-1 D326K | 5 nM, 5 nM | SyxLE/Syb | 2.5 nM, 5 nM | 9.4 ± 0.2 |
| Munc18-1 WT | 5 nM, 5 nM | SyxLE/Syb | 2.5 nM, 5 nM | 17.9 ± 1.0 |
| Munc18-1 D326K | 30 nM, 35 nM | SyxLE M183A/Syb | 15 nM, 35 nM | n.d.[b] |
| Munc18-1 D326K | 30 nM, 35 nM | SyxLE D184P/Syb | 15 nM, 35 nM | n.d.[b] |
| Munc18-1 P335A | 5 nM, 5 nM | SyxLE/Syb | 2.5 nM, 5 nM | 23.6 ± 4.0 |
| Munc18-1 S42Q | 35 nM, 30 nM | SyxLE/Syb | 35 nM, 15 nM | 267 ± 28 |
| Munc18-1 L307R | 20 nM, 20 nM | SyxLE/Syb | 10 nM, 20 nM | 72.2 ± 1.7 |
| Munc18-1 Q301D | 20 nM, 20 nM | SyxLE/Syb | 10 nM, 20 nM | 90.6 ± 5.2 |
| Munc18-1 E352K | 10 nM, 20 nM | SyxLE/Syb | 10 nM, 10 nM | 97.8 ± 5.5 |
| Munc18-1 L348R | 20 nM, 30 nM | SyxLE/Syb | 20 nM, 15 nM | 140 ± 137 |

[a]Six experiments were performed for each pair of proteins, 3 at the first concentrations indicated in the [Protein 1] and [Protein 2] columns, and 3 at the second concentrations listed. The six $K_D$ values obtained where used to calculated the listed average $K_D$ ± standard deviation.

[d]Binding was too weak to derive accurate $K_D$s (n.d. = not determined)
